# LARGE, an AMPA receptor interactor, plays a large role in long-term memory formation by driving homeostatic scaling-down

**DOI:** 10.1101/236547

**Authors:** Bo Am Seo, Taesup Cho, Daniel Z. Lee, Hwa Young Lee, Joong-Jae Lee, Boyoung Lee, Seong-Wook Kim, Kathryn A. Cunningham, Kelly T. Dineley, Thomas A. Green, Ho Min Kim, Se-Young Choi, Hee-Sup Shin, Myoung-Goo Kang

## Abstract

Dynamic trafficking of AMPA-type glutamate receptor (AMPA-R) in neuronal cells is a key cellular mechanism for learning and memory in the brain, which is regulated by AMPA-R interacting proteins. LARGE, a protein associated with intellectual disability, was found to be a novel component of the AMPA-R protein complex in our proteomic study. Here, our functional study of LARGE showed that during homeostatic scaling-down, increased LARGE expression at the Golgi apparatus (Golgi) negatively controlled AMPA-R trafficking from the Golgi to the plasma membrane, leading to downregulated surface and synaptic AMPA-R targeting. In *LARGE* knockdown mice, long-term potentiation (LTP) was occluded by synaptic AMPA-R overloading, resulting in impaired long-term memory formation. These findings indicate that the fine-tuning of AMPA-R trafficking by LARGE at the Golgi is critical for memory stability in the brain. Our study thus provides novel insights into the pathophysiology of brain disorders associated with intellectual disability.

## Introduction

*LARGE* is expressed strongly in the brain (particularly the hippocampus), relative to other tissues(**`Peyrard et al., 1999**). In humans, mutations in *LARGE* are associated with congenital muscular dystrophy type 1D, which is characterized by clinical features including profound intellectual disability, abnormal electroretinogram findings, and subtle structural brain abnormalities (**Clarke et al., 2011; Longman et al., 2003; Vaillend et al., 2008; Lisi and Cohn, 2007**). *Large^myd^* mice, which carry a natural truncation mutation of *LARGE,* exhibit a number of neurological phenotypes, including sensorineural deafness and defective retinal transmission, along with developmental brain abnormalities (**Holzfeind et al., 2002; Michele et al., 2002**) and impaired long-term potentiation (LTP) (**Satz et al., 2010**). These human and mouse studies suggest that abnormal synaptic function may be responsible for intellectual disabilities in human patients with *LARGE* mutations.

In our previous proteomic analysis, we found that LARGE forms a protein complex with the AMPA-type glutamate receptor (AMPA-R) (**Kang et al., 2012**). Excitatory glutamatergic synaptic transmission within the central nervous system is primarily mediated by AMPA-R, as well as NMDA-type glutamate receptor (NMDA-R), and increasing numbers of proteins have been found to form complexes with and thus regulate the dynamic trafficking of AMPA-R. This tight regulation of AMPA-R trafficking in and out of the synapses, mediated by AMPA-R interactors, is widely considered to be a central brain mechanism involved in information storage (**Hanley, 2010**).

Changes in neuronal activity can alter synaptic transmission efficacy. This phenomenon, known as synaptic plasticity, is a main mechanism underlying learning and memory in the brain. Several forms of synaptic plasticity, including Hebbian, homeostatic, and structural, have been identified. Hebbian synaptic plasticity involves acute adaptations of neurons in the brain during learning and memory processes (including LTP), wherein repeated neuronal stimulation causes changes in synaptic efficacy. Homeostatic synaptic plasticity involves the chronic adaptation of neurons against prolonged changes in neuronal activity and is required for stability of encoded memory in the brain. Structural synaptic plasticity describes changes in the lengths, shapes, and numbers of neuronal dendrites and dendritic spines. Notably, AMPA-R trafficking plays critical roles in all three types of synaptic plasticity. Therefore, our understanding of synaptic plasticity, learning and memory, and cognitive brain function relies on knowledge about the molecular mechanisms underlying AMPA-R trafficking regulation (**Huganir and Nicoll, 2013**).

Our functional study of LARGE revealed a novel and robust cellular mechanism underlying AMPA-R trafficking from the Golgi to the cell surface, which contributes to all three types of synaptic plasticity. LARGE is necessary for synaptic scaling-down, a type of homeostatic plasticity. Specifically, it downregulates the synaptic targeting of AMPA-R by negatively modulating AMPA-R trafficking from the Golgi to the cell surface. Synaptic AMPA-R overloading due to LARGE deficiency causes hippocampal LTP occlusion and the abnormal enlargement of dendritic spines, resulting in abrogated long-term memory formation. LARGE thus contributes to the stability of encoded memory by fine-tuning AMPA-R trafficking at the Golgi in the hippocampal neurons.

## Results

### LARGE is necessary for neuronal homeostatic scaling-down

Previous studies of LARGE have suggested a role for this protein in synaptic plasticity (**Clarke et al., 2011; Holzfeind et al., 2002; Lisi and Cohn, 2007; Longman et al., 2003; Michele et al., 2002; Peyrard et al., 1999; Satz et al., 2010; Vaillend et al., 2008**). Our proteomic study identified LARGE as a component of the AMPA-R protein complex (**Kang et al., 2012**), a major player in synaptic plasticity via dynamic trafficking in and out of the neuronal surface and synapse. To determine whether LARGE could regulate AMPA-R trafficking, we selected a homeostatic scaling method that would allow the monitoring of surface and synaptic AMPA-R trafficking in response to changes in neuronal activity (**Turrigiano, 2008**).

First, we monitored LARGE protein expression in cultured hippocampal neurons treated with either tetradotoxin (TTX) or bicuculline for 48 h to induce homeostatic scaling-up and scaling-down, respectively (**Turrigiano, 2008**), which were confirmed by monitoring changes in GluA1 surface localization (i.e., expected increases and decreases in response to TTX or bicuculline treatment, respectively) through cell surface biotinylation. Interestingly, bicuculline treatment led to a significant increase in LARGE expression, whereas TTX had no effect on LARGE (**Figure 1A and Figure 1-figure supplement 2A**). We more carefully analyzed this increase in LARGE expression from 0 to 72 h after bicuculline treatment and, excitingly, observed an inverse correlation between LARGE and surface GluA1/2 expression (**Figure 1B**). These results strongly suggest an association between increased LARGE expression and decreased AMPA-R surface localization. To test this possibility, we altered LARGE expression using adeno-associated virus (AAV) expressing short hairpin (sh) RNA and rescue constructs (**Figure 1C**) after validating the efficacy of shRNA-mediated *LARGE* KD (**Figure 1-figure supplement 1**). Notably, the bicuculline-induced decrease in surface GluA1 was mitigated by *LARGE* KD but reversed by *LARGE* rescue (**Figure. 1C**). However, neither LARGE KD nor LARGE rescue affected the TTX-induced increase in surface GluA1 (**Figure 1-figure supplement 2B**).

**Figure 1.**
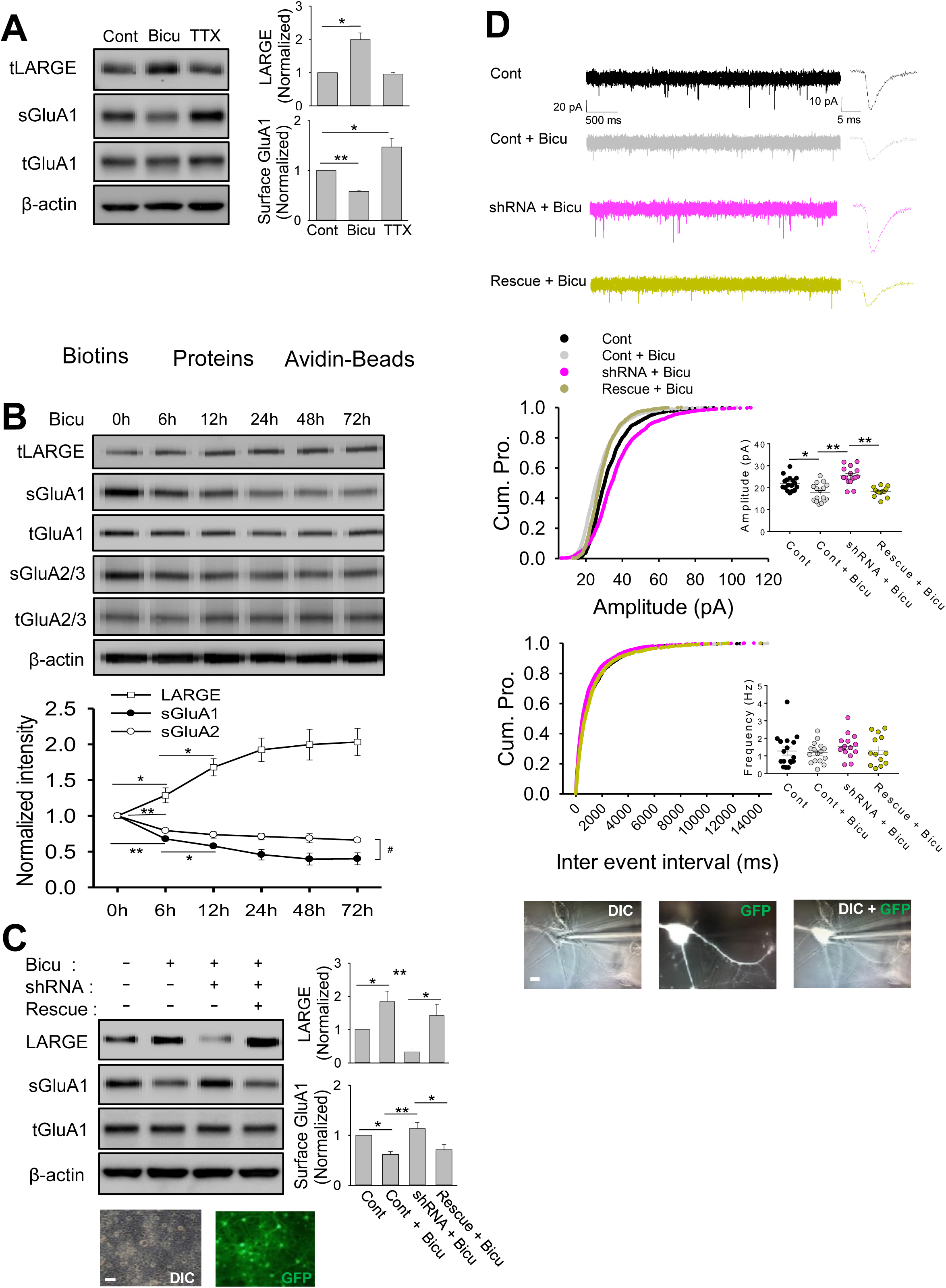
LARGE is necessary for neuronal homeostatic scaling-down (**A**) Total LARGE expression increased significantly during synaptic scaling-down in response to a 2-day bicuculline treatment (Bicu), but was not significantly affected by 2-day tetradotoxin (TTX)-induced synaptic scaling-up. Our surface biotinylation approach (schema) revealed increased and decreased surface GluA1 expression levels in response to treatment with TTX and Bicu, respectively, which confirmed the respective induction of scaling-up and - down (n = 4; two-tailed t-test, \**P* <0.01,\*\**P* <0.001). (**B**) During synaptic scaling-down, inverse correlations of LARGE expression were observed with surface GluA1 and GluA2 expression (n = 6; one-way ANOVA, \**P* <0.05, \*\**P* <0.005). The decrease of surface GluA1 was greater than that of GluA2 (n = 6; two-way RM ANOVA, #P <0.001). (**C**) High-density hippocampal cultures were infected with an AAV expressing scrambled shRNA with GFP (Cont), *LARGE* shRNA with GFP (shRNA) or *LARGE* rescue (Rescue). At DIV (Day of *in vitro)* 14, the neurons were subjected to surface biotinylation after a 2-day Bicu or TTX treatment. The Bicu-induced decrease in surface GluA1 expression was blocked by the knockdown (KD) of endogenous LARGE with *LARGE* shRNA (shRNA), but reversed by *LARGE* rescue (Rescue) (n = 4; one-way ANOVA, \**P* <0.05, \*\**P* <0.005). (**D**) The Bicu-induced decrease in mEPSC amplitude was blocked by the shRNA-mediated knockdown (KD) of endogenous *LARGE,* but was reversed by *LARGE* rescue (Rescue) (1300, 1300, 1300, and 1262 events from n = 17, 16, 15, and 13 neurons, respectively; one-way ANOVA; amplitude, F(3,57) =15.558, \**P* <0.005, \*\**P* <0.001; frequency; F(3,57) =0.619, *P* =0.606). Representative images show GFP-positive and negative neurons in a whole-cell patch clamp experiment. Cultured hippocampal neurons transfected on DIV (Day of *in vitro)* 8 with cDNA plasmids expressing either scrambled shRNA with GFP, *LARGE* shRNA with GFP, or *LARGE* shRNA with GFP and LARGE rescue with GFP, were subjected to a whole-cell patch clamp experiment after a 2-day treatment with Bicu or TTX. A calcium phosphate method was used to yield a transfection efficiency of approximately 20%. Scale bar = 10 μm.

Next, we monitored changes in synaptic AMPA-R in a single cell level by measuring the miniature excitatory postsynaptic current (mEPSC) at 48 h after bicuculline or TTX treatment with or without *LARGE* shRNA or rescue (**Figure 1D and Figure 1-figure supplement 2C**). Here, *LARGE* KD mitigated the bicuculline-induced decrease in synaptic AMPA-R according to changes in mEPSC amplitude, and *LARGE* rescue reversed this occlusion (**Figure 2D**). Again, neither *LARGE* KD nor *LARGE* rescue affected the TTX-induced increase in synaptic AMPA-R (**Figure 1-figure supplement 2C**). The fidelity of our mEPSC experiments was verified by monitoring cell batch-to-batch variation (**Figure 1-figure supplement 3**).

**Figure 2.**
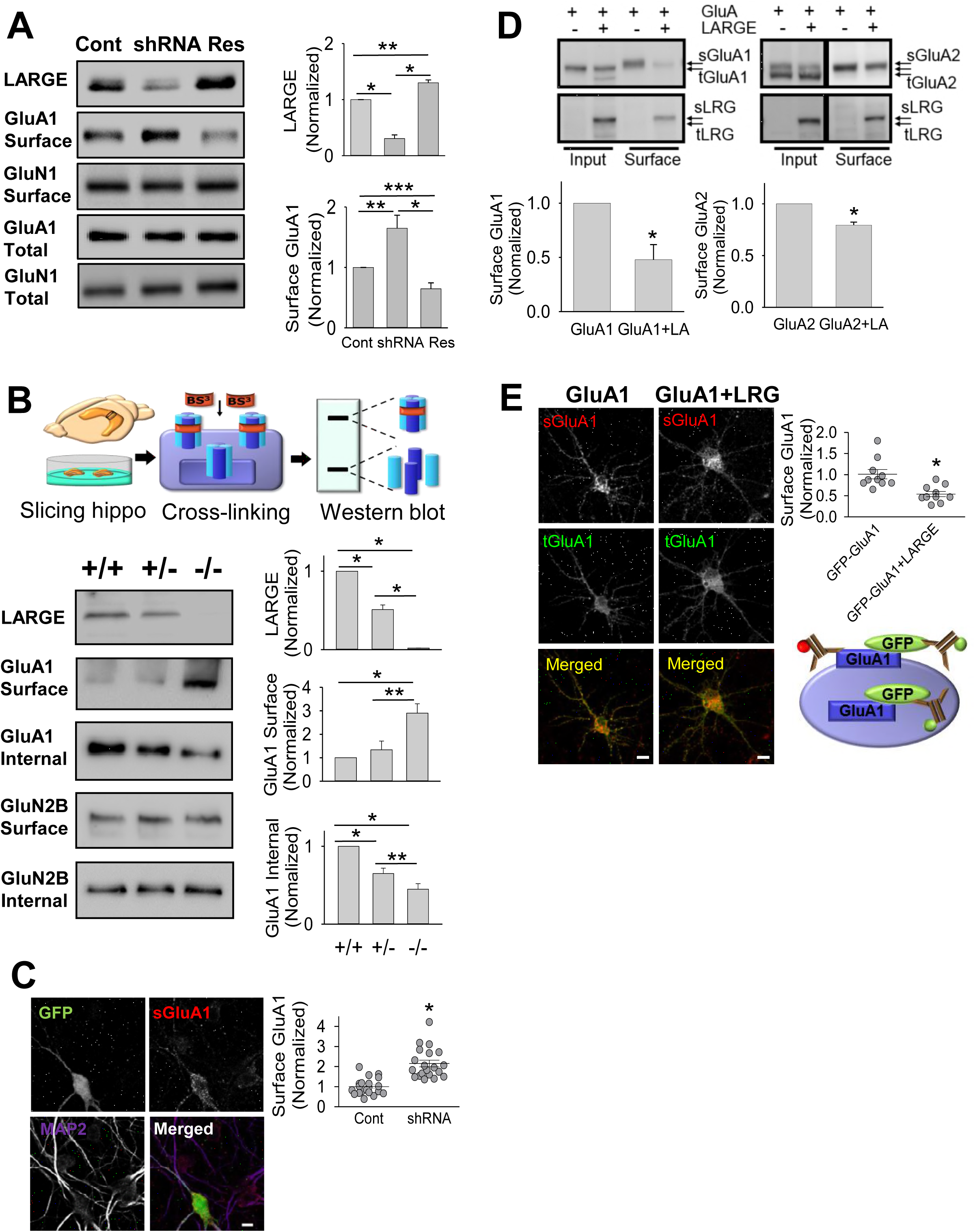
LARGE downregulates the surface localization of AMPA-R. (**A**) *LARGE* knockdown (KD) increased cell-surface GluA1 (Surface) expression, which was reversed by *LARGE* rescue. However, changes in LARGE expression did not affect the total GluA1 (Input) or GluN1 expression (n = 3; one-way ANOVA, \**P* <0.001, \*\**P* <0.005, \*\*\**P* <0.05). (**B**) A schema of the *ex vivo* cross-linking of surface proteins using BS^3^. Cross-linked and non-cross-linked proteins are indicated by high-molecular-weight (Surface) and low-molecular-weight bands (Internal), respectively. Surface GluA1 levels increased and internal GluA1 levels decreased in homozygous (-/-) mice (Large^myd^) *ex vivo,* compared with wildtype (+/+) and heterozygous (+/-) mice. GluN2B was used as an internal control (n = 3; twotailed t-test, \**P* <0.001, \*\**P* <0.05). (**C**) An imaging analysis of cultured neurons revealed higher levels of surface-localized GluA1 (sGluA1) in neurons expressing *LARGE* shRNA (GFP-positive neurons) than in neighboring GFP-negative neurons (Cont). The GluA1 antibody was applied before neuron permeabilization. MAP2, a marker of neuronal dendrites, was used to determine the number and morphology of neurons (n = 20, 20 neurons; MannWhitney rank-sum test, T = 234, U = 24, \**P* <0.001). Scale bars = 10 μm. (**D**) Co-expression of LARGE-GFP (LARGE or LRG) with myc-GluA1 (GluA1) and myc-GluA2 (GluA2) in HEK293T cells significantly reduced the respective cell-surface localization of GluA1 (sGluA1) and GluA2 (sGluA2). However, total GluA1 (tGluA1) and GluA2 (tGluA2) levels in the Input were not significantly altered (n = 8; two-tailed t-test, \**P* <0.001). (E) Co-expression of LARGE with GFP-GluA1 significantly reduced cell-surface GluA1 (sGluA1) levels at cultured neurons. A schema of our imaging assay. sGluA1 was labeled with a red fluorophore-conjugated anti-GluA1 antibody prior to neuron permeabilization. After permeabilization, total GluA1 (tGluA1) was stained using a green fluorophore-conjugated anti-GFP antibody (n = 10, 10 neurons; Mann-Whitney rank-sum test, T = 147, U = 8, \**P* <0.005). Scale bar = 10 μm.

These data strongly suggest that LARGE is necessary for homeostatic scaling-down in hippocampal neurons.

### LARGE downregulates AMPA-R surface localization

LARGE is required for AMPA-R surface and synaptic targeting during homeostatic scaling-down (**Figure 1**). The inverse correlation between LARGE and surface GluA1/2 expression (**Figure 1B**) suggests a link between increased LARGE expression and decreased AMPA-R surface localization, and recent studies have indicated that AMPA-R synaptic targeting is mainly regulated by the abundance of surface AMPA-R (**Granger et al., 2012; Hanley, 2010**). Accordingly, we hypothesized that LARGE drives synaptic scaling-down by downregulating AMPA-R surface targeting, and we first investigated whether LARGE could downregulate the cell-surface localization of AMPA-R (**Figure 2**).

Our surface biotinylation approach demonstrated significant increases and decreases in AMPA-R surface localization in response to *LARGE* KD and rescue, respectively, whereas KD or rescue affected total AMPA-R expression or NMDA-R surface localization (**Figure. 2A**). We additionally evaluated AMPA-R surface localization *ex vivo* using membrane-impermeable bis(sulfosuccinimidyl) suberate (BS^3^), which crosslinks proteins exposed on the cell surface (**Figure 2B**). The molecular weight of cross-linked surface GluA1 (>250 kDa) was higher than that of intracellular GluA1 (~105 kDa), consistent with our previous study (**Lee et al., 2012**). Although the surface and intracellular GluA1 levels respectively increased and decreased significantly in *Large^myd-/-^* mice relative to wild-type mice, NMDA-R surface localization was not affected by LARGE deficiency (**Figure 2B**), further confirming that LARGE specifically modulates AMPA-R. The increased AMPA-R surface localization following *LARGE* KD was originated in a single cell level (**Figure 2C**).

We further investigated whether LARGE could modulate AMPA-R surface localization in heterologous expression system. Here, in HEK293T cell culture, LARGE overexpression led to strong decreases in surface GluA1 and GluA2 localization but did not affect total AMPA-R expression (**Figure 2D**). The observed greater decrease in GluA1 relative to GluA2 (**Figure 2D**) may explain the greater decrease in surface GluA1 vs. GluA2 during synaptic scaling-down (**Figure 1B**). Again, the effect of LARGE overexpression on AMPA-R surface localization was recapitulated in individual hippocampal neurons (**Figure 2E**).

### LARGE upregulates the Golgi localization of AMPA-R

Next, we used confocal imaging to assess the subcellular localization of LARGE and thus understand the mechanism by which LARGE downregulates AMPA-R surface targeting. LARGE is known to localize at the Golgi in heterologous cells (**Brockington et al., 2005**) and to function at the Golgi in myocytes (**Kanagawa et al., 2004**). Similarly, in cultured hippocampal neurons, we observed a major pool of LARGE at the Golgi and Golgi outposts (**Figure 3A**). Accordingly, we hypothesized that LARGE downregulates AMPA-R trafficking from the Golgi to cell surface by increasing AMPA-R localization at the Golgi.

**Figure 3.**
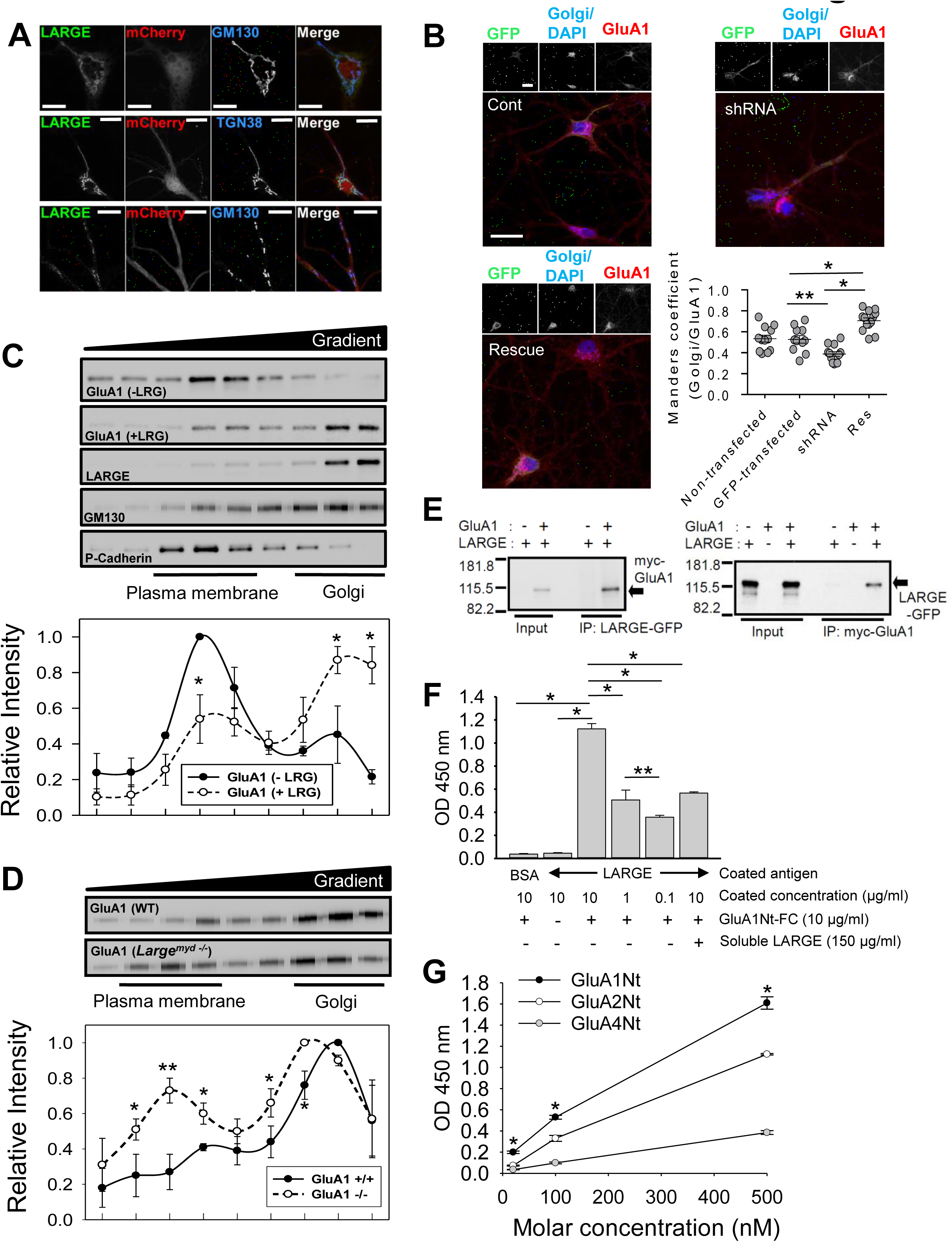
LARGE directly interacted with AMPA-R to increase AMPA-R localization at the Golgi. (**A**) Confocal images demonstrating major pools of GFP-tagged LARGE (LARGE-GFP) in the cis-Golgi (Top), trans-Golgi (middle), and Golgi outposts (bottom) of cultured hippocampal neurons. GM130 and TGN38 are cis-Golgi and trans-Golgi marker proteins, respectively. Scale bar = 10 μm. (**B**) *LARGE* knockdown (KD; shRNA) decreased the number of GluA1 molecules at the Golgi, compared with non-transfected and scrambled-shRNA transfected neurons (Cont). This phenomenon was reversed by *LARGE* rescue (Rescue) (n = 12, 11, 12, 13 neurons; F(2,33) = 31.618, \**P* <0.001, \*\**P* <0.002). Scale bar = 30 μm. (**C**) The density gradient-based subcellular fractionation of HEK293T cells revealed that the co-expression of LARGE with GluA1 (+LRG) significantly altered the distribution of AMPA-R pools in the Golgi and plasma membrane. In the +LRG group, the relative size of the GluA1 pool at the Golgi (GM130) increased significantly whereas that in the plasma membrane (P-Cadherin) decreased significantly relative to the control group (-LRG, GluA1 only) (n = 3; two-tailed t-test, \**P* <0.05). (**D**) Subcellular fractionation of hippocampal tissues from *Large* knockout mice (Large^myd-/-^) revealed relative decreases and increases, respectively, in the relative sizes of the GluA1 pools in the Golgi and plasma membrane, compared with those in wild-type mice (WT) (n = 3, 3 mice; two-tailed t-test, \**P* <0.05, \*\**P* <0.001). (**E**) Heterologous HEK293T cells in which LARGE-GFP had been co-expressed with myc-tagged GluA1 (myc-GluA1) were subjected to immunoprecipitation (IP). IP of LARGE with an anti-GFP antibody specifically co-immunoprecipitated GluA1 (left), and IP of GluA1 with an anti-myc antibody specifically co-immunoprecipitated LARGE (right). (**F**) The direct interaction between LARGE and AMPA-R was examined using an enzyme-linked immunosorbent assay (ELISA). The LARGE ectodomain (Catalytic domain 1 [CD1] + CD2) used to coat the bottom of the plate bound directly to the Fc-fused N-terminal ectodomain of GluA1 (GluA1-Fc). Binding was quantified by measuring the activity of FC antibody-coupled peroxidase (n = 3; F(5,12) = 293.046, \**P* <0.001, \*\**P* <0.01). (**G**) Another ELISA used to evaluate the relative binding affinities of LARGE for each AMPA-R subunit yielded the following order from highest to lowest: GluA1 > GluA2 > GluA4 (n = 3; F(2,12) = 867.644, \**P* <0.001 compared with GluA2 and GluA4). (**B, F**) One-way ANOVA, (**G**) two-way RM ANOVA.

To test this concept, we first analyzed the co-localization of GluA1 with the Golgi marker GM130 in cultured hippocampal neurons after manipulating LARGE expression. Confocal imaging demonstrated that *LARGE* KD significantly decreased the pool of GluA1 at the Golgi, whereas *LARGE* rescue reversed this phenomenon (**Figure 3B**). Similarly, in HEK293T cells, LARGE overexpression significantly increased AMPA-R localization at the Golgi (**Figure 3-figure supplement 1A**).

Next, we biochemically analyzed the effects of LARGE co-expression on subcellular AMPA-R localization by fractionating subcellular organelles from HEK293T cells transfected with GluA1 without (-LRG) or with LARGE (+LRG) (**Figure 3C**). Without LARGE coexpression, the major GluA1 pool was detected in fractions enriched for P-cadherin, a plasma membrane marker. With LARGE co-expression, however, the major GluA1 pool shifted to high-density fractions enriched for GM130, a Golgi marker. Furthermore, the distributions of GluA1 and LARGE in the gradient almost completely overlapped when the proteins were co-expressed (**Figure 3C**), indicating a strong and direct association. Moreover, the relative AMPA-R pool size in the Golgi fractions decreased significantly in the brains of *LARGE* KO mice relative to wild-type mice, whereas the relative pool size in the plasma membrane fractions increased significantly (**Figure 3D and Figure 3-figure supplement 1B**). These results suggest that LARGE plays an important role in maintaining AMPA-R pools at the Golgi.

### LARGE associates with AMPA-R through direct interactions

HEK293T cells do not express synaptic proteins. Therefore, the co-sedimentation and colocalization of GluA1 with LARGE in these cells (**Figure 3C** and **Figure 3-figure supplement 1A**) suggested a direct interaction. To test this possibility, we performed reciprocal co-immunoprecipitation (co-IP) of LARGE and AMPA-R from HEK293T cells. Both co-IP strategies consistently showed that LARGE could specifically bind to AMPA-R in non-neuronal cells that do not express other known AMPA-R-binding proteins (**Figure 3E**). Interestingly, in a co-IP of LARGE with GluA2, LARGE bind only to the GluA2 correspond to intracellular GluA2 (**Hall et al., 1997**) (**Figure 3-figure supplement 2A**), suggesting that LARGE binds with AMPA-R within the cell, likely at the Golgi.

We further reconstituted the interaction of LARGE with AMPA-R *in vitro* using an enzyme-linked immunosorbent assay (ELISA) (**Figure 3F,G**). Given the molecular structures and topologies of LARGE and GluA1, we hypothesized that the C-terminal ectodomain of LARGE would bind the N-terminal ectodomain of GluA1. After confirming the purification of each protein (**Figure 3-figure supplement 2B-E**), we constructed an ELISA assay to verify the specific and direct interaction of LARGE with GluA1 (**Figure 3F**). Notably, GluA1 bound to LARGE with a higher affinity relative to that exhibited by GluA2 or GluA4 (**Figure 3G**). The interactions of LARGE with GluA1, 2, and 4 indicated that LARGE binding to AMPA-R is not subunit-specific. Similarly, the binding of the other AMPA-R interacting proteins, such as Stargazin (**Tomita et al., 2003**) and CKAMP44 is also not subunit-specific (**von Engelhardt et al., 2010**).

As GluA1 binds directly to the LARGE ectodomains (including catalytic domains), we evaluated whether the interaction of LARGE with AMPA-R could affect glycosylation of the latter. However, we observed no dramatic changes in LARGE-mediated GluA1 glycosylation (**Figure 3-figure supplement 3**), suggesting that the physical interaction of LARGE with GluA1, rather than LARGE glycosyltransferase activity, is the essential element with respect to AMPA-R. Similarly, a previous report found that the physical interaction of Notch with OFUT1, a glycosyltransferase, rather than OFUT1 enzymatic activity, was essential to the role of OFUT1 in Notch trafficking (**Okajima et al., 2005**). These results (**Figure 3**) consistently demonstrate the ability of LARGE to interact directly and specifically with AMPA-R to increase localization of the latter protein at the Golgi.

### The pool of LARGE-associated AMPA-R at the Golgi increases during homeostatic scaling-down

Next, we investigated whether LARGE-interacting AMPA-R at the Golgi might increase during homeostatic scaling-down. Confocal imaging of cultured hippocampal neurons revealed a significant increase in LARGE (**Figure 4A**) and accompanying increase in GluA1 at the Golgi (**Figure 4B**) at 48 h after bicuculline treatment, which also led to an increase in GluA1 and LARGE co-localization around the perinuclear areas of neurons (**Figure 4C**). Together, these data strongly suggest that the co-localization of GluA1 and LARGE at the Golgi increases during synaptic scaling-down. Indeed, subcellular fractionation confirmed increases in both GluA1 and LARGE at the Golgi during scaling-down (**Figure 4D**). Finally, co-IP confirmed a significant increase in the association of LARGE with GluA1 during bicuculline-induced synaptic scaling-down (**Figure 4E**). Although the amount of GluA1 immunoprecipitated by a GluA1 antibody decreased after bicuculline treatment, probably because of increased protein (e.g., LARGE) binding and consequently reduced epitope exposure, the amount of LARGE that co-immunoprecipitated with GluA1 remained significantly elevated. Together, our results strongly support our working model, wherein increased LARGE expression (in response to increased neuronal activity) negatively regulates AMPA-R trafficking from the Golgi to the plasma membrane, thus downregulating AMPA-R synaptic targeting during synaptic scaling-down (**Figure 4F**).

**Figure 4.**
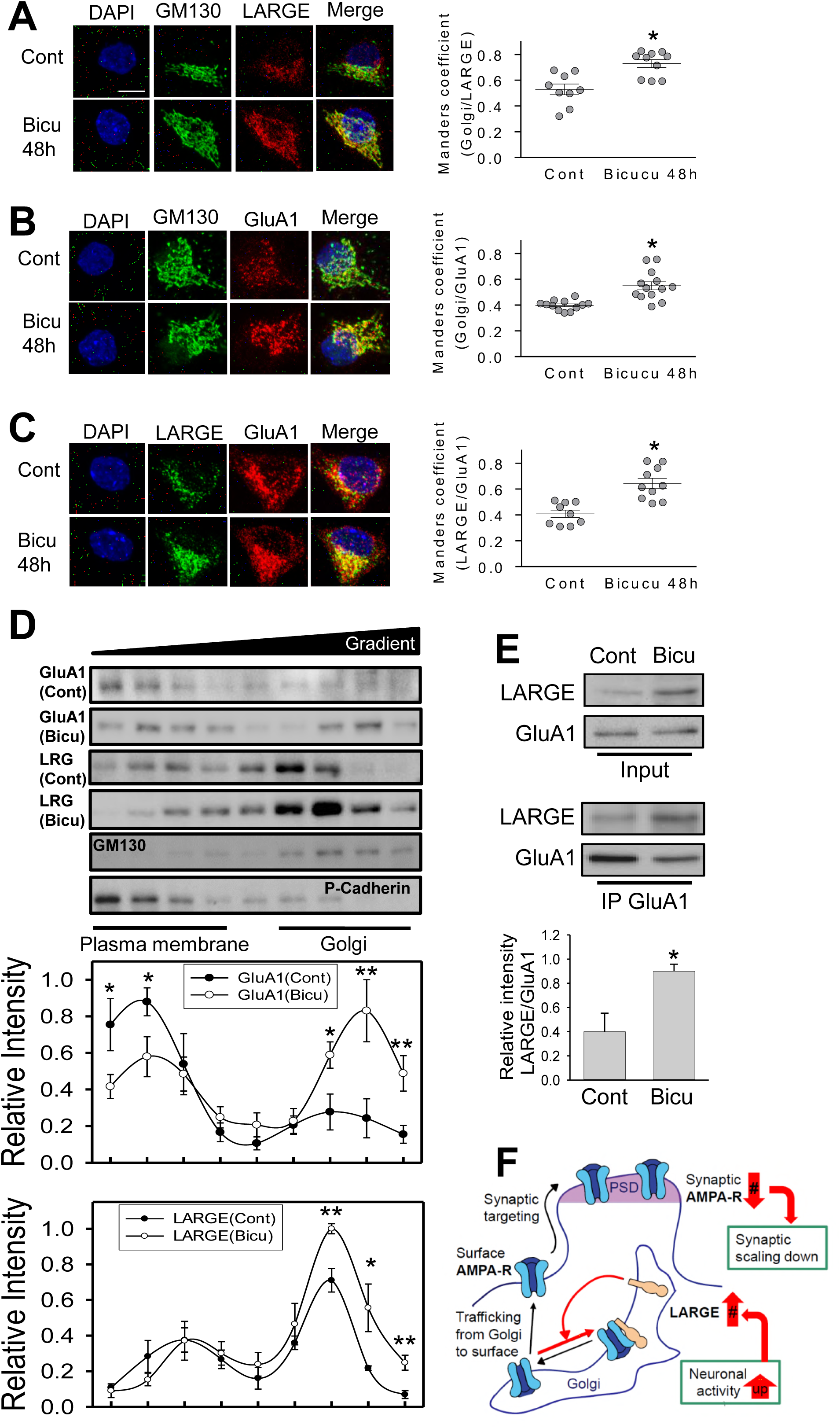
The LARGE-associated pool of AMPA-R at the Golgi increases during homeostatic scaling-down. (**A-C**) A series of confocal microscopy images of cultured hippocampal neurons double-stained for a Golgi marker (GM130) and LARGE or GluA1 yielded several findings. (**A**) LARGE localization at the Golgi increased significantly after a 48-h bicuculline treatment (Bicu 48h) (n = 9, 10 neurons; two-tailed t-test, t_17_ = −3.937, \**P* <0.001). Scale bar = 10 μm. (**B**) GluA1 localization at the Golgi increased significantly in response to Bicu 48h (n = 12, 13 neurons; Mann-Whitney rank-sum test, T = 90, U = 12, \**P* <0.001). (**C**) The colocalization of GluA1 and LARGE in the perinuclear region increased significantly in response to Bicu 48h (n = 9, 10 neurons; two-tailed t-test, t_17_ = −4.728, \**P* <0.001). (**D**) Density gradient fractionation of subcellular organelles revealed significant increases in the relative amounts of GluA1 and LARGE in the Golgi fraction following treatment with Bicu 48h (n = 4; two-tailed t-test, \**P* <0.05, \*\**P* <0.005). (**E**) Bicu 48h significantly increased the binding of LARGE to GluA1, as demonstrated by the increased co-immunoprecipitation of LARGE and GluA1 (n = 3; two-tailed t-test, \**P* <0.01). (**F**) Schema of our working model for the regulation of AMPA-R trafficking by LARGE during homeostatic scaling-down.

### *LARGE* KD impairs hippocampal LTP due to synaptic AMPA-R overload

Previous studies of LARGE have suggested a role for this protein in Hebbian synaptic plasticity (**Clarke et al., 2011; Holzfeind et al., 2002; Lisi and Cohn, 2007; Longman et al., 2003; Michele et al., 2002; Peyrard et al., 1999; Satz et al., 2010; Vaillend et al., 2008**), including LTP (**Satz et al., 2010**). The LTP deficit in *LARGE* KO mice could be due to abnormal brain development such as neuronal migration defect (**Holzfeind et al., 2002; Satz et al., 2010**). To determine whether the LARGE could affect Hebbian synaptic plasticity in normally developed adult mouse brain, we investigated hippocampal synaptic plasticity via *in vivo* LTP after KD of LARGE after the stereotaxic injection of an AAV expressing *LARGE* shRNA with GFP. Theta-patterned stimulation (TPS) protocol (**Cho et al., 2013**) readily induced long-lasting hippocampal CA1 LTP in control mice, but not in *LARGE* KD mice (**Figure 5A**). Despite this LTP impairment, *LARGE* KD exhibited dramatic increases in field excitatory postsynaptic potential (fEPSP) amplitudes (**Figure 5A**), leading us to analyze these amplitudes at different stimulation intensities. In an input-output analysis, the AMPA-R fEPSP amplitudes in *LARGE* KD mice increased significantly over input intensities of 30–100 mV (). However, *LARGE* KD did not affect presynaptic neurotransmitter release and short-term plasticity, which were evaluated using the paired-pulse ratio (PPR). The PPRs at inter-pulse intervals of 200, 100, 75, 50, and 25 ms did not differ between control and *LARGE* KD mice (**Figure 5-figure supplementary 1B**), suggesting that the synaptic changes observed in the latter mice are not presynaptic events.

**Figure 5.**
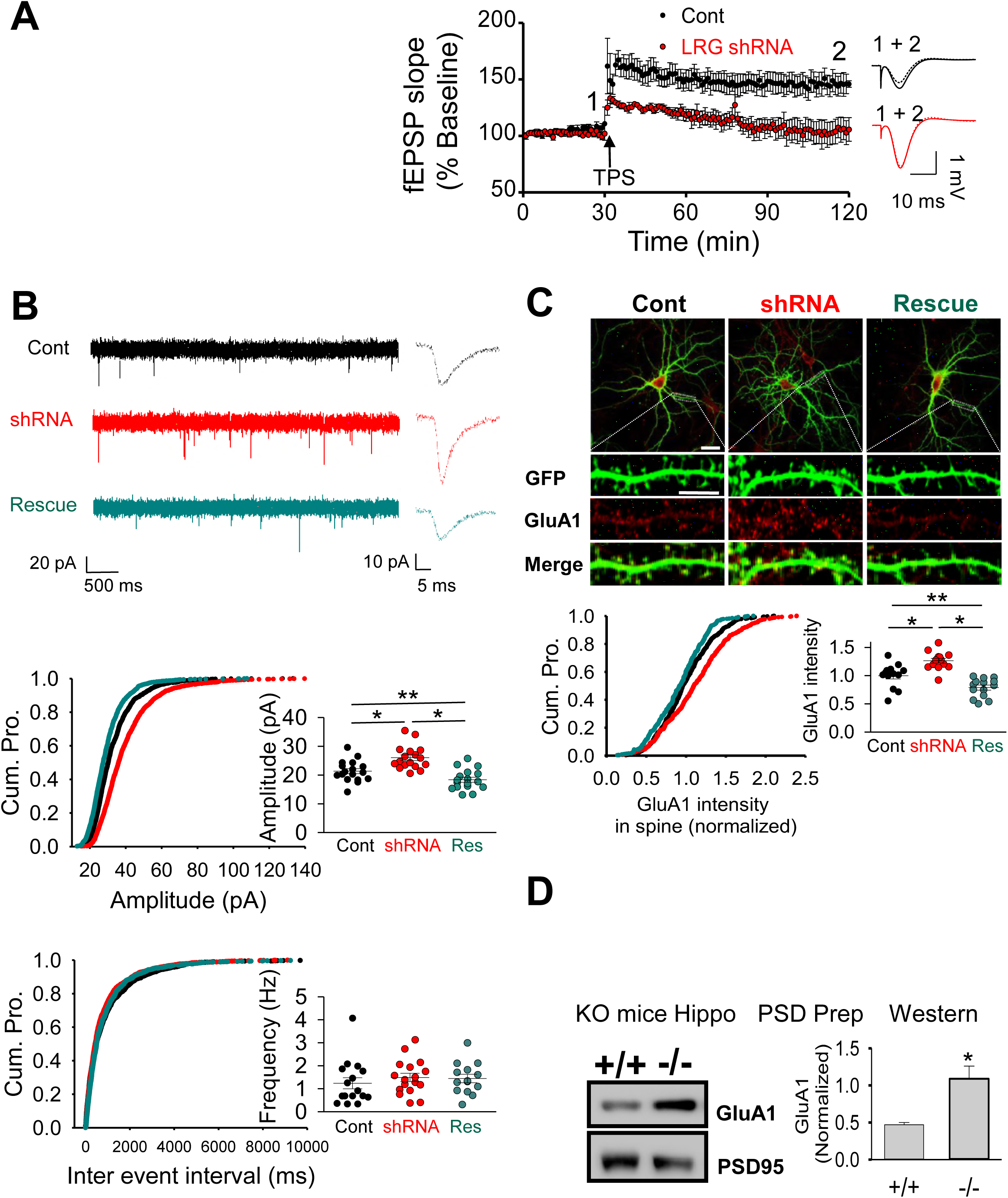
*LARGE* knockdown (KD) causes synaptic AMPA-R overload and thus impairs longterm potentiation (LTP) in the hippocampal CA1 region. (**A**) A schema, plot, and traces from an *in vivo* LTP analysis after an injection of virus encoding *LARGE* shRNA. *LARGE* KD (LRG shRNA) was found to impair LTP (n = 8, 8 mice; two-tailed t-test, \**P* <0.05). Scale bar = 300 μm. (**B**) Whole-cell patch clamping yielded current traces, cumulative plots, and scattered plots from an mEPSC analysis following transfection with plasmids encoding scrambled shRNA (Cont), *LARGE* shRNA, or *LARGE* rescue. *LARGE* KD increased the amplitude but not the frequency of mEPSC, whereas *LARGE* rescue reversed this amplitude change. Three different groups from 48 coverslips in four batches of neuronal culture were recorded (n = 1653, 1660, and 1622 events from *n =* 17, 17, and 17 neurons, respectively; Amplitude, F(_2_,4_8_) = 17.815, \**P* <0.001, **P = 0.026; Frequency, F_(2_,_48_) = 0.452, *P* = 0.639; Cont: black, shRNA: red, Rescue: cyan). (**C**) *LARGE* KD (shRNA) increased the number of GluA1 molecules in the dendritic spines, whereas this phenomenon was reversed by *LARGE* rescue (n = 529, 529, and 562 spines from *n =* 13, 14, and 14 neurons, respectively; F(2,38) = 25.026, \**P* <0.001, \*\**P* <0.005). Scale bars = 30 μm (whole cell image) and 10 μm (dendrite). (**D**) Schema, blot, and bar graph of a western blot analysis of GluA1 expression in the postsynaptic density (PSD) of hippocampi from *LARGE* knockout (KO) mice (-/-), demonstrating increased synaptic AMPA-R expression relative to that observed in WT mice (+/+) *in vivo (n =* 3, 3 mice; two-tailed t-test, \**P* <0.005). (b, c) one-way ANOVA.

The above results strongly suggest that *LARGE* KD increases the synaptic current by increasing the number of AMPA-R molecules at the postsynapses. We therefore further examined whether LARGE could regulate the synaptic localization of AMPA-R. First, we analyzed AMPA-R-mediated mEPSC in cultured neurons transfected with scrambled shRNA, *LARGE* shRNA, or a *LARGE* rescue plasmid. *LARGE* KD significantly increased the amplitude, but not the frequency, of mEPSC relative to the controls. Moreover, *LARGE* rescue completely reversed the effect of KD on amplitude (**Figure 5B**). However, when we tested the effect of *LARGE* KD on inhibitory synapses, we observed no change in the miniature inhibitory postsynaptic current (mIPSC) in either *LARGE* KD or rescue cells (**Figure 5-figure supplementary 1C**), consistent with the findings of a previous study(**Pribiag et al., 2014**). These mEPSC and mIPSC analyses, therefore, demonstrate that changes in LARGE expression specifically affect the number of synaptic AMPA-R molecules at the excitatory postsynapses, without affecting presynapses or inhibitory synapses.

Second, we subjected cultured hippocampal neurons to confocal imaging to demonstrate that the number of GluA1 molecules within the dendritic spine increased significantly with *LARGE* KD relative to control neurons, and this increase was reversed by *LARGE* rescue (**Figure 5C**). Moreover, *LARGE* KD neurons had significantly larger spine heads but similar spine densities relative to control neurons; again, this was reversed by *LARGE* rescue (**Figure 5-figure supplementary 2**). Finally, our biochemical analysis demonstrated that GluA1 expression in the postsynaptic density (PSD) increased significantly in the hippocampi of *LARGE* KO mice relative to controls (**Figure 5D**). Altogether, the impaired hippocampal LTP (**Figure 5A**) observed with *LARGE* KD is probably attributable to synaptic AMPA-R overload (**Figure 5B-D**), which inhibits the further capacity to increase the synaptic AMPA-R pool during LTP.

### LARGE deficiency impairs fear memory

The effects of LARGE KD on Hebbian (LTP) and structural (spine size) synaptic plasticity (**Figure 5**), which underlie learning and memory in the brain, strongly suggested a role for LARGE in cognitive functions in the brain. To determine whether LARGE deficiency could cause learning and memory problems in animals, we subjected *LARGE* KO *(Large^myd-/-^),* wild-type, and heterozygous mice *(Large^myd+/-^* and *+^/-^)* to Pavlovian fear conditioning (**Figure 6A**) and observed similar freezing behaviors in all three groups (**Figure 6B**). One day after fear conditioning, the mice were subjected to tests of contextual memory, an index of associative memory dependent on both hippocampal and amygdala function, and cued memory, a hippocampus-independent index of associative memory that still relies on proper amygdala function (**Sanders et al., 2003**). Compared with wild-type mice, KO mice displayed significant reductions in freezing behavior during both contextual and cued memory tests (**Figure 6C,D**), indicating that *LARGE* KO leads to deficits in both hippocampus- and amygdala-dependent memory.

**Figure 6.**
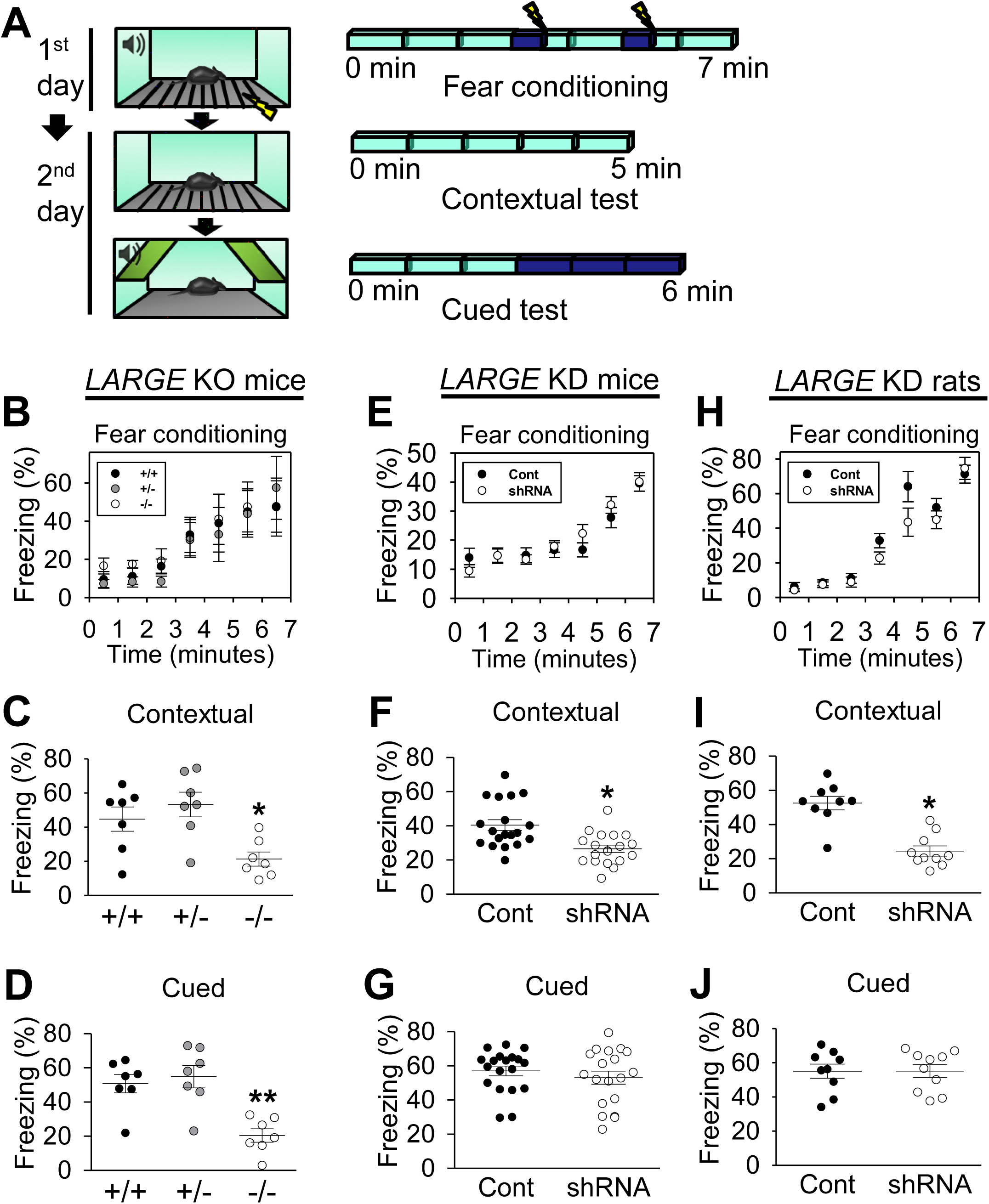
LARGE deficiency impairs fear memory. (**A**) Schema of a fear memory test. (**B**) *LARGE* knockout (KO) did not induce differences in fear conditioning (n = 7, 7, 7 mice) (**C, D**) Contextual and cued fear memory deficits were observed in KO mice (-/-). (n = 7, 7, 7 mice; contextual, F(2,18) = 6.808, \**P* <0.01; cued, F(2,18) = 12.148, \*\**P* <0.001). (**E**) No differences in fear conditioning were observed in *LARGE* knockdown (KD) mice (n = 19, 19 mice). (**F**) Contextual memory deficits were observed in *LARGE* KD mice (n = 19, 19 mice, two-tailed t-test, t36 = 3.684, \**P* <0.01). (**G**) No differences were observed in cued memory (n = 19, 19 mice, Mann-Whitney rank-sum test, T = 390, U = 161, *P* = 0.579). (**H**) No differences in fear conditioning were observed in *LARGE* KD rats (n = 9, 10 mice). (**I**) Contextual memory deficits were observed in *LARGE* KD rats (n = 9, 10 rats, two-tailed t-test, t_17_ = 5.709, \**P* <0.005). (**J**) No differences were observed in cued memory (n = 9, 10 mice, two-tailed t-test, t_17_ = −0.0162, *P* = 0.987). (**E-J**) Animals were injected with a virus expressing either scrambled (Cont) or *LARGE* shRNA with GFP (shRNA). (**B, E, H**) Twoway RM ANOVA, (**C, D**) One-way ANOVA.

We note, however, that KO mice are constitutive mutants. Therefore, the observed memory deficits may be attributable to abnormal brain development. Accordingly, we knocked down *LARGE* in the bilateral hippocampal CA1 regions of adult mice and rats via the stereotaxic injection of an AAV expressing *LARGE* shRNA with GFP prior to Pavlovian fear conditioning to examine the potential effects of LARGE on memory processes in the absence of life-long inherent developmental abnormalities. Another group of mice injected with AAV expressing scrambled shRNA with GFP served as a control. We subsequently validated the efficacy of shRNA-mediated *LARGE* KD *in vivo* (**Figure 6-figure supplementary 1**) and confirmed the reliability of the experimental animals used in fear tests (**Figure 6-figure supplementary 2**). Although the two groups exhibited similar freezing behavior (**Figure 6E**), *LARGE* KD mice exhibited significantly less freezing behavior during contextual but not cued memory tests (**Figure 6F,G**). In rats, *LARGE* knockdown in the hippocampal CA1 produced the same effects on memory (**Figure 6H-J**).

### LARGE deficiency impairs hippocampus-dependent long-term memory

We next subjected *LARGE* KD mice to various memory tests to identify the specific memory-associated role of LARGE in the brain. An initial open field test revealed no significant differences between KD and control groups (**Figure 7A**). In other words, *LARGE* KD mice exhibit normal locomotion and anxiety levels. To test spatial working memory, we used a Y-maze to evaluate whether KD mice could remember which arms of the maze had been recently visited. Again, no significant differences were observed between the groups (**Figure 7B**).

**Figure 7.**
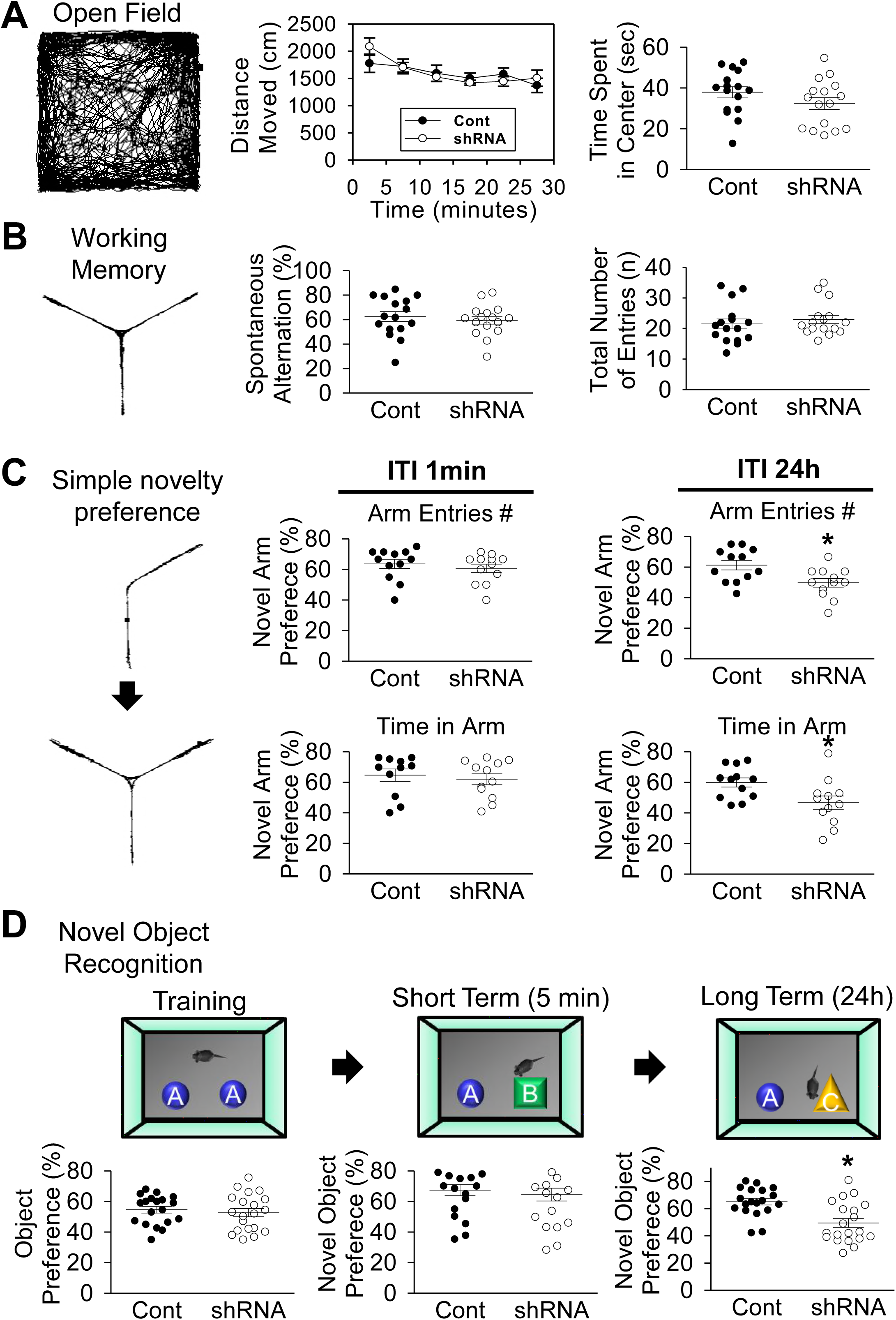
LARGE deficiency impairs hippocampus-dependent long-term but not short-term memory. (**A**) In the open field test, no differences were observed in the distances moved and time spent in the center (n = 16, 16 mice; center time, two-tailed t-test, t_30_ = 1.393, *P* = 0.174). (**B**) In the Y-maze-based working memory test, no differences were observed in spontaneous alternations or the total number of entries (n = 16, 16 mice, two-tailed t-test, t_30_ = 0.581, *P* = 0.566). (**C**) In the simple novelty preference test, *LARGE* knockdown (KD) mice exhibited a preference in the short term (1 min; n = 12, 12 mice, Mann-Whitney rank-sum test; entries, T = 168, U = 54, *P* = 0.308; time, T = 141, U = 57, *P* = 0.601), but not in the long term (24 h; n = 12, 12 mice, two-tailed t-test; entries, t_22_ = 2.727, \**P* = 0.012; time, t_22_ = 2.483, **P = 0.021). ITI: inter-trial intervals. The novel arm preference was calculated as (novel/novel + other) x 100%. (**D**) In the novel object recognition test, control and *LARGE* KD mice similarly explored two identical objects during training (n = 19, 20 mice, two-tailed t-test, t_37_ = 0.556, *P* = 0.582). Both groups exhibited similar preferences during a short-term memory test (5 min; n = 19, 20 mice, Mann-Whitney rank-sum test, T = 393, U = 177, *P* = 0.725). In a long-term memory test, *LARGE* KD mice exhibited no preference for the novel object (24 h; n = 19, 20 mice, two-tailed t-test, t_37_ = 3.773, \**P* <0.001). The novel object preference was calculated as (novel/novel + familiar) x 100%. (**A**) Two-way RM ANOVA for distance moved.

Next, we used a simple novelty preference test that has used in previous studies of GluA1 function for memory (**Freudenberg et al., 2016; Sanderson et al., 2009**), to measure both short- and long-term spatial memory in *LARGE* KD mice, using inter-trial intervals of 1 min and 24 h, respectively (i.e., hippocampus-dependent memory) (**Sanderson et al., 2009**). Although both groups exhibited a similar degree of preference for the novel arm at 1 min, the *LARGE* KD group failed to exhibit a preference for the novel arm at 24 h, compared with the control group (**Figure 7C**).

Finally, we performed the novel object recognition test, a popular testing paradigm for hippocampal function as a relay point of recognition memory (**Stilling et al., 2014**). During the training session, both groups displayed similar degrees of preference for two equal objects, with no inter-group difference in the short-term (5 min) preference for a novel object. Over the long term (24 h), however, the *LARGE* KD group failed to display a preference for the novel object relative to the control group (**Figure 7D**). Taken together, these findings suggest that hippocampal *LARGE* KD specifically impairs long-term, but not short-term, spatial and recognition memory.

## Discussion

LARGE downregulates AMPA-R synaptic targeting by negatively controlling AMPA-R trafficking from the Golgi to the plasma membrane, and thus fine-tunes synaptic AMPA-R abundance (**Figure 4F**), a novel cellular mechanism underlying homeostatic scaling-down, which is a form of homeostatic plasticity. Moreover, the fine-tuning of AMPA-R trafficking by LARGE contributes to the two other types of synaptic plasticity, Hebbian (LTP) and structural (spine size) synaptic plasticity, that are the main underlying mechanisms of learning and memory. Indeed, hippocampus-dependent long-term memory formation was impaired in LARGE knockdown mice. *LARGE* mutations are associated with intellectual disabilities in humans (**Clarke et al., 2011; Longman et al., 2003; Vaillend et al., 2008**), and abnormal homeostatic synaptic scaling has been suggested as a pathophysiological component of brain disorders associated with maladaptive synaptic plasticity(**Turrigiano, 2008**). Our study thus provides novel insights into psychiatric and neurological disorders associated with intellectual disability.

### LARGE function for synaptic plasticity and memory stability

During homeostatic scaling-down, an increased LARGE expression at the Golgi apparatus negatively modulated AMPA-R trafficking from the Golgi to the plasma membrane, leading to the downregulation of surface and synaptic AMPA-R targeting. This novel mechanism for the regulation of AMPA-R trafficking via Golgi explains how global synaptic scaling-down at the single cell level is possible with all synapses regardless of their locations, connections, and histories.

Both Hebbian and homeostatic synaptic plasticity mainly adjust synaptic strength by altering the abundance of AMPA-R in the postsynaptic membrane. Although the common output suggests crosstalk between homeostatic and Hebbian synaptic plasticity (**Turrigiano, 2008**), the mechanism of interaction within the same neuron remained unclear. The emerging idea that homeostatic synaptic plasticity acts as a form of metaplasticity to influence the subsequent induction of Hebbian plasticity (**Arendt et al., 2013; Soares et al., 2013**) is supported by our study. The lack of scaling-down and consequent increased GluA1 synaptic targeting precluded the further synaptic addition of AMPA-R required for the induction of LTP. Moreover, our study strongly suggested that the crosstalk is required to stabilize encoded memories in the brain. A memory can be stabilized in the long-term through a consolidation process. LARGE KD mice exhibit deficits not in short-term memory but in long-term memory that needs consolidation (**Figure 7**). Without the LARGE-mediated scaling-down, synaptic AMPA-R levels increase chronically. The AMPA-R overload at the synapse is similar to an unconstrained LTP status (**Turrigiano, 2008**), wherein memory remains unstable because of a breakdown in synapse specificity. Together, our LARGE functional study proposed a novel mechanism underlying the role of homeostatic synaptic plasticity for memory stability.

### Function of LARGE in the hippocampal consolidation of long-term memory

The fear memory deficit observed in *LARGE* KO and KD animals indicates a role for LARGE in hippocampus-dependent memory (**Figure 6**). To determine a more specific function of LARGE, we subjected hippocampal CA1 *LARGE* KD mice to working, spatial, and recognition memory analyses. Working memory, which is necessary for the temporary storage and manipulation of information required for complex cognitive tasks, involves the hippocampal CA1 region (**Dillon et al., 2008**). The absence of *LARGE* KD-induced changes in working memory (**Figure 7B**) indicated that LARGE affects a particular type of memory. Next, we applied simple novelty preference and novel objective recognition tests to respectively examine spatial and recognition memory in *LARGE* KD mice. The former test identified an impairment in long-but not short-term spatial memory (**Figure 7C**). In simple novelty preference test, spatial long-term memory is formed through repetitive training over several days. This result thus suggests that LARGE plays a role in hippocampal memory consolidation. Similarly, the novel objective recognition test revealed impairment in long-rather than short-term object recognition in *LARGE* KD mice (**Figure 7D**). Working memory is hippocampus-dependent (**Dillon et al., 2008**) but does not require consolidation (**Guitar and Roberts, 2015**). Hence, the intact spatial working memory and short-term memory observed in this study strongly suggest that hippocampal *LARGE* KD specifically affects hippocampus-dependent, consolidation-requiring processes essential for memory stability.

### Effects of LARGE on Hebbian synaptic plasticity

Hippocampal *LARGE* KD caused a failure of LTP (**Figure 5A**), which underlies long-term memory formation (including spatial memory consolidation) (**Lynch, 2004; Nabavi et al., 2014**). This LTP impairment was attributed to an overload of synaptic AMPA-R that blocked further AMPA-R synaptic targeting (**Figure 5B-D and Figure 5-figure supplementary 1A**). In consistent with our results, LTP occlusion due to elevation in postsynaptic AMPAR surface expression and function was reported previously (**Traunmuller et al., 2016**). The synaptic AMPA-R overload was due to chronic increase of neuronal activity (**Figure 4F**). Supportively, previous studies have shown that abnormally increased synaptic activity impairs LTP and memory. For example, enhanced synaptic responses suppress LTP development, resulting in hippocampal memory deficits(**Barnes et al., 1994**), whereas chronically increased fEPSPs cause persistent deficits in the acquisition of new spatial information (**Castro et al., 1989; McNaughton et al., 1986**). However, short-term plasticity remained intact in *LARGE* KD animals (**Figure 5-figure supplementary 1B**), consistent with the normal acquisition of fear training and short-term memory in both KO and KD animals (**Figure 6,7**). These results corroborate the theory that short- and long-term memories result from dissociable physiological processes (**Spear and Miller, 1981**) and are formed by different neurobiological mechanisms(**Barker et al., 2006**). The lack of change in short-term plasticity also supports our hypothesis that LARGE has a postsynaptic, rather than presynaptic, effect on synaptic plasticity, as demonstrated by the absence of effects of *LARGE* KD or overexpression on mEPSC frequency (**Figure 5B**).

Studies of LARGE function in the brain have focused on the association of LARGE with the dystrophin glycoprotein complex (DGC) (**Michele et al., 2002; Pribiag et al., 2014; Satz et al., 2010**). DGC components, including dystroglycan, are specifically expressed in inhibitory GABAergic synapses but not at excitatory glutamatergic synapses (**Levi et al., 2002; Pribiag et al., 2014**). Accordingly, the effects of LARGE on Hebbian synaptic plasticity via the regulation of AMP-R abundance at excitatory synapses provides novel insights into the intellectual disabilities associated with *LARGE* mutations in the brain.

## Materials and Methods

### Animal care and treatments

Experiments involving animals were performed in accordance with procedures approved by the Institutional Animal Care and Use Committee at the University of Texas Medical Branch (UTMB) and the Korea Advanced Institute of Science and Technology (KAIST). Care was taken to minimize the number of animals used and their discomfort. Colonies of *Large^myd^* mice originated from Jackson Laboratory and were transferred from Dr. Kevin Campbell’s laboratory at the University of Iowa prior to establishment at UTMB. Male adult C57BL/6 mice (wild type) were used in this study. The Sprague-Dawley rats and C57BL/6 mice used for behavior tests and neuronal cell culture were purchased from Harlan (USA) or Orient (Korea).

### HEK293T cell culture

HEK293T cells were cultured at 37°C in high-glucose Dulbecco’s Modified Eagle’s Medium (Sigma) supplemented with 10% fetal bovine serum (Gibco) and antibiotics (Gibco). For transfection, cells were plated on coverslips in 12-well plates or 6-well plates and then transfected with various expression plasmids using the transfection reagent TransIT-X2 (Mirus #mir6000).

### Neuronal culture

Hippocampal neuronal cultures were prepared and maintained with glia-conditioned media as previously described (Kang et al., 2009). Briefly, timed pregnant female C57BL/6 mice and Sprague-Dawley rats were purchased, and primary cells were isolated from embryonic day 17-18 (E17-18) pups. Mixed cell cultures containing both neurons and glia were then grown on coverslips in 12-well plates and in 6-well plates for use in biochemical experiments. The cultures were treated with 5 μM cytosine β-D-arabinofuranoside (AraC, Sigma) at day in vitro (DIV) 3 to reduce the number of glial cells.

### Preparation of cDNA, shRNA, virus, transfection

The LARGE cDNA plasmid was kindly provided by Dr. Kevin Campbell (University of Iowa, Iowa City, IA, USA) and has been described previously (Kanagawa et al., 2004). The shRNA virus constructs were designed, generated, and screened as previously described (Kang et al., 2009). Briefly, four siRNAs were designed using Custom *SMARTpool* Design and synthesized (GE Dharmacon). The knockdown efficacy of each siRNA was tested, and the sequence of the selected siRNA (rat: CGGCUUUGCUGCCUUGAAA, mouse:UGGCUUUGCUGCCUUGAAA) was used to design an shRNA. The *LARGE* rescue construct was generated by replacing the sequence encoding GFP in the pAAV vector used for *LARGE* knockdown with a *LARGE* cDNA containing a shRNA-resistant sequence (CGGCUUUGCUGCCUUGAAA => CGGCUUUGCUGCCCUGAAA).

The AAV was packaged and purified as follows. The shRNA designed from the selected siRNA sequence was subcloned into the pAAV vector, and subsequently packaged into the virus by co-transfecting HEK293T cells with pHelper and pAAV-RC (serotype DJ/8). At 72 h post-transfection, the viral particles were harvested through two freeze/thaw cycles and sonication. Benzonase and Rnase I were added to the virus-released solution. To remove cell debris, the cell lysates were centrifuged at 2500 χ g for 15 min, and the supernatants were filtered through 0.2-μm syringe filters (Millipore, USA). A stock solution of 2.5 N NaCl and 40% PEG8000 (Sigma #5413) was added to the supernatant to yield final respective concentrations of 0.5 N and 8%. The resulting solution was incubated on ice for 3 h and centrifuged at 2000 χ g for 30 min, after which the supernatant was discarded. The pellets containing AAV were resuspended in HEPES buffer, and this crude AAV solution was treated with chloroform and PEG for an aqueous two-phase extraction (10% PEG8000-13.2% (NH4)2SO4) and final dialysis. The titer (>1 χ 10^11^ TU/ml) was measured by treating 10^6^ neurons with the AAV and measuring enhanced green fluorescent protein (GFP) expression after 1 week (Figures S1A). The knockdown efficacy of AAV was evaluated using western blotting (Figures S1A).

Cell transfection procedures used the same shRNA constructs (pAAV) except for the rescue construct. For these experiments, the *LARGE* rescue construct was generated by adding a self-cleaving 2A peptide (P2A) site between the sequence encoding the *LARGE* cDNA containing the shRNA-resistant sequence and enhanced GFP within the *LARGE* shRNA-containing pAAV vector. The knockdown and rescue efficacies were evaluated using confocal imaging.

### Reverse transcription polymerase chain reaction (RT-PCR)

RNAs were extracted (MACHERY-NAGEL # MN740955.50) from the infected hippocampal CA1. cDNAs were synthesized from these RNAs using a kit according to the manufacturer’s protocol (Invitrogen #11904-018). The genes *(LARGE, β-actin)* tested in this study were amplified from cDNA using DNA polymerase (enzynomics #P525) and gene-specific primers (in Key resource table).

### Stereotaxic injection of virus into animals

Adult (9-10 weeks of age) male C57BL/6 mice and Sprague-Dawley rats (body weight: 275-300 g) were assigned to either a control or an experimental group and injected with AAV expressing either scrambled or *LARGE* shRNA, respectively, with enhanced GFP. Specifically, the animals were anesthetized with 2% avertin (Sigma) and placed into a stereotaxic apparatus (David Kopf instruments), after which the virus solution was injected into the bilateral CA1 region of the hippocampus using the following coordinates: mouse, anteroposterior, −1.95 mm from bregma; mediolateral, ±1.39 mm; dorsoventral −1.66 mm; rat, anteroposterior, −2.4 mm from bregma; mediolateral, ± 2.0 mm; dorsoventral −2.0 mm. For mice, 0.5 μl of virus solution was injected using a *picospritzer* with a glass pipette (diameter, 15-20 μm). For rats, 2 μl of virus solution was injected at a rate of 0.1 μl/min using a syringe pump with a glass pipette (5-μl syringe), and the needle was left in place for at least 10 min post-infusion. The animals were allowed to recover in a heated chamber before waking and were used for behavior tests at least 3 weeks after virus injection. After the tests, the targeting of virus injection into the hippocampal CA1 region was confirmed by the digital imaging of brain slices (2 mm thick) under a blue LED flashlight (DFP-1; NightSea) to excite the GFP. After detecting GFP fluorescence, the fluorescent region of the hippocampus was dissected for biochemical analyses.

### Fear conditioning and fear memory tests

Fear conditioning and fear memory tests were performed as previously described (Hernandez et al., 2010). The two-pair model of fear conditioning involves placing the animal in the fear conditioning apparatus (Med Associates) for a total of 7 min. Animals were left to explore freely for 3 min. At the 3-min and 5-min time points, an acoustic conditioned stimulus (white noise, 80 dB) was delivered for 30 s, and an unconditioned footshock stimulus was administered through the grid floor during the last 2 s of tone presentation (0.5-0.6 mA for mice, 0.8 for rats) and co-terminated with the tone. Contextual fear memory was evaluated 24 h after paired training by placing the animal into the same training context and measuring freezing behavior for 5 min. The cued fear memory was evaluated at least 4 h after the contextual test by placing the animal in a different context (novel cage floor, lighting, odor, and visual cues) with a 3-min free exploration period. At the 3-min mark, the same acoustic conditioned stimulus was delivered for 3 min, and freezing behavior was measured using Actimetrics FreezeFrame software with real-time digital video. Data are expressed as the percentage of freezing during each minute or as a mean across all minutes.

### Measurement of shock threshold

The shock thresholds for flinching, jumping, and vocalization, which are used as indices of sensitivity to a shock stimulus, were measured as previously described (Hernandez et al., 2010). Each animal was placed in the fear conditioning apparatus, and a sequence of single foot shocks was delivered. Initially, a 0.1-mA shock was delivered for 1 s; thereafter, the shock intensity was increased by 0.1 mA at 30-s intervals until an intensity of 1.0 mA was reached. The shock intensity was then decreased by 0.1 mA at 30-s intervals until an intensity of 0.1 mA was reached. Thresholds were then quantified by averaging the shock intensity at which each animal gave a flinching, jumping, and vocalization response.

### Open field test

In mice, locomotor activity and anxiety-related behavior were measured using the open field test. A mouse was placed in the corner of an open field box (40 cm × 40 cm × 40 cm; material, white acryl). To evaluate locomotor activity, the total distance traveled (cm) was measured during each 5 min of the 30-min test. To evaluate anxiety, the time spent in the center of the box during the first 5 min (s) was analyzed (Ethovision XT 8.5, Nodulus). A photometer was used to adjust the light within a range of 5-10 lux.

### Y-maze

A Y-maze comprising of three symmetrical arms at 120° angles (30 cm length × 12 cm height × 7 cm width) was constructed from opaque acryl. A mouse was placed in the center of the maze and allowed to freely explore the three arms for 5 min. Timing began once the mouse left the center. Arm entry was defined as having all four limbs inside an arm. The sequence of entries was recorded to calculate spontaneous alternations.

### Simple novelty preference test

The simple novelty preference test was performed as described previously (Sanderson et al., 2009). The mice received five 2-min training trials involving exposure to two arms of a Y-maze (Start and Other arms; the third arm is blocked). Short and long-term memory were assessed by changing the interval between exposure training sessions. Timing was started once the mouse left the start arm. After exposure training, mice were subjected to a novelty preference test in which they were allowed to explore all three arms of the maze (Novel, Start, and Other arms) for 2 min. The exposure trials and novelty preference test were each separated by either 1 min (1-min inter-trial interval [ITI] condition) or 24 h (24-h ITI condition). The novel arm preference was calculated as the percentage ratio of the total amount of time spent exploring and number of entries into the Novel and Other arms (Novel/Novel + Other) x 100%.

### Novel object recognition test

The novel object preference test was performed as described previously (Stilling et al., 2014). Mice were habituated to a white acryl box for 5 min on each of 2 consecutive days. Mice were then habituated to the same two objects placed in corners of the box for 5 min on each of the 2 consecutive days. The following day, the objects were exchanged for two new, identical objects (A + A), and the mice were allowed to explore the objects for 5 min. Next, the mice were placed in their home cages for 5 min (short-term memory task) and re-exposed to the arena in which one object had been exchanged (A + B). After 24 h, B was exchanged for C (long-term memory task). The durations of object contacts were measured. The novel object preference was reported as: (novel / sum (both objects)) x 100%.

### *In vivo* field EPSP recordings

Three to 4 weeks prior to *in vivo* field recording, AAV expressing *LARGE* shRNA with GFP was infused into the hippocampal CA1 regions of adult C57BL/6 mice (8-9 weeks old) as described in *Stereotaxic injection of virus into animals.* The stereotaxic unilateral injection of 0.5 μl of higher-titer AAV (1 x 10^11^ TU/ml) was performed using a stereotaxic, motorized nano-injector (World Precision Instruments) at a rate of 0.1 μl/min via a Hamilton syringe connected to a microinjection pump.

The fEPSPs from the hippocampal CA1 region were recorded as previously described (Cho et al., 2013). C57BL/6 mice (*n* = 11-12) were anesthetized with urethane (1.6 g/kg, i.p.; Sigma) and placed into a stereotaxic frame. Rectal temperature was maintained intraoperatively at 36.5°C ± 0.5°C using a temperature controller (Harvard Instruments). The scalp was opened and separated. Trephine holes were drilled into the skull, and electrodes were positioned in the area of the hippocampal stratum radiatum. A bipolar stimulating electrode (2.0 mm posterior to bregma, 2.0 mm lateral to midline) was used for Schaffer collateral stimulation, and a monopolar recording electrode (1.9 mm posterior to bregma, 1.4 mm lateral to midline) was used to record from the CA1 region. The final depths of the electrodes were adjusted to optimize the magnitude of the evoked responses. The fEPSPs were adjusted to 50-60% of the maximal response size for testing. Stimulation was applied using an analog-to-digital interface (1322A; Molecular Devices) and a Digital Stimulus Isolation unit (Getting Instruments). The pyramidal neuron responses to Schaffer collateral stimulation were recorded using a differential amplifier (P55 A. C. pre-amplifier; Grass Instruments) and analyzed using WinLTP software (WinLTP Ltd.). Responses were evoked by single-pulse stimuli delivered at 20-s intervals. A stable baseline was recorded for 30 min. LTP was induced by applying theta-patterned stimulation (TPS, four trains comprising of 10 bursts of five pulses at 400 Hz with a 200-ms inter-burst interval and a 20s inter-trial interval) to the CA1 and was optimized based on previous studies (Cho et al., 2013).

### Whole-cell patch-clamp recordings for mEPSCs and mIPSC analyses

Cultured hippocampal neurons were prepared as described above. Using a calcium phosphate transfection kit (Invitrogen #K278001), cultured hippocampal neurons were transfected at DIV 8 with cDNA plasmids expressing either scrambled shRNA with enhanced GFP, *LARGE* shRNA with GFP, or *LAEGE* shRNA with GFP and LARGE rescue. The cultured neurons were used for electrophysiological recordings at 3 days post-transfection. Miniature excitatory postsynaptic currents (mEPSCs) and miniature inhibitory postsynaptic currents (mIPSCs) were recorded at room temperature (21-23°C). Whole-cell voltage-clamp recordings were performed using a multiclamp 700B amplifier (Molecular Devices), filtered at 1 KHz, and digitized at 10 KHz (Digidata 1550; Molecular Devices). Recording pipettes (4-6 MΩ) were filled with the following intracellular solutions, as appropriate: for mEPSC analysis, 140 mM Cs-MeSO4, 8 mM NaCl, 10 mM HEPES, 0.5 mM EGTA, 1 mM MgCl2, 4 mM Mg-ATP, 0.4 mM Na-GTP, 5 mM QX-314; for mIPSC analysis, 130 mM CsCl, 10 mM NaCl, 1.1 mM EGTA, 2 mM MgCl_2_, 0.1 mM CaCl_2_, 10 mM HEPES, 2 mM Mg-ATP. The pH was adjusted to 7.2 using CsOH, with 280-290 mOsm.

Hippocampal neurons on coverslips were transferred to a recording chamber that was continuously perfused with extracellular solution (pH 7.4, 310-320 mOsm) containing 150 mM NaCl, 3.1 mM KCl, 2 mM CaCl_2_, 1 mM MgCl_2_, 10 mM HEPES, and 25 mM glucose. One micromolar tetrodotoxin (Tocris Bioscience #1078), 50 μM DL-AP5 (Tocris Bioscience #3693), and 100 μM picrotoxin (Sigma #P1675) were always included in the extracellular perfusing solution for mEPSC (for mIPSC, 20 μM CNQX (Tocris Bioscience #0190) was used instead of picrotoxin). All recordings were voltage clamped at −70 mV. Acquired data were analyzed using pCLAMP 10.6 (Molecular Devices). Access resistance was continuously monitored. The data were discarded if the *R_a_* varied by >20% during recording. Changes in frequency and amplitude were analyzed, quantified, and presented using traces, cumulative plots, and scatter plots. We confirmed that the frequency and amplitude did not vary among the batches used in experiments or between non-transfected and GFP-transfected neurons (Cont). For the homeostatic synaptic scaling experiment, neurons were pre-incubated in either 1 μM TTX or 20 μM bicucullin for 48 h before mEPSC recordings were obtained as described above.

### Preparation of PSD

PSD was prepared as described previously (Kang et al., 2012), with some modifications.

### Surface biotinylation

HEK293T cells in 10-cm plates or cultured neurons in 6-well plates were washed with phosphate-buffered saline (PBS) or artificial cerebrospinal fluid (ACSF; 0.15 M NaCl, 10 mM HEPES, 3 mM KCl, 0.2 mM CaCl_2_ dihydrate, 10 mM glucose), respectively, and placed on ice. The cells were then incubated with 1-1.5 mg/ml sulfo-NHS-SS-biotin (Thermo #21331) for 20 min at 10°C. Subsequently, the biotin was quenched by incubation with 50 mM glycine for 10 min on ice, followed by washing. The cells were removed by scraping and solubilized in IP buffer containing 1.0% Triton X-100 and 0.1% SDS for 1 h at 4°C. After solubilization, the cells were centrifuged at 14,000 x g for 15 min, and the pellet was discarded. The supernatant containing biotinylated proteins was incubated with NeutrAvidin-Sepharose beads (Thermo #29200) for 3 h at 4°C. The beads were then thoroughly washed with IP buffer, and the proteins were eluted for 5 min at room temperature (RT) with protein gel-loading buffer. The total and cell-surface proteins were analyzed by SDS-PAGE, followed by western blot analysis.

### Analysis of cell-surface proteins in hippocampal slices using the cross-linking reagent BS^3^

Cell-surface proteins were labeled using the membrane-impermeable cross-linking reagent bis(sulfosuccinimidyl) suberate (BS^3^, Thermo #21585) as described in previous studies (Lee et al., 2012), with some modifications. After the cardiac perfusion of mice with cold ACSF, the hippocampi were removed and placed into cold ACSF oxygenated with a 95% O2/5% CO2 gas mixture. Each hippocampus was cut into five ~1-mm-thick slices and allowed to float in oxygenated ACSF. Ten slices from two hippocampi were incubated in BS^3^ (Thermo-Pierce) solution (2.5 mM BS^3^ dissolved in ACSF) for 40 min at 10°C with gentle shaking. After quenching with ACSF containing 100 mM glycine three times for 5 min each, the slices were processed for solubilization, SDS-PAGE, and western blotting as described above for surface biotinylation.

### Fractionation of subcellular organelles from cells or tissues

The iodixanol-based iso-osmotic density gradient-based subcellular organelle fractionation procedure was developed according to the instructions of the kit manufacturer (OptiPrep, Sigma #D1556), and optimized for two scales (5-ml volume gradient and 0.7-ml volume gradient). Briefly, cells or brain lysates were centrifuged for 10 min at 3000 x g, and the pellet was discarded. The supernatant was then centrifuged for 1 h at 100,000 x g to remove cytosolic contamination. The resulting second pellet was applied to the top of an OptiPrep discontinuous iodixanol gradient formed by the stepwise addition of solutions with increasing percentages of iodixanol (diluted in PBS). The 5.0-ml volume gradient was formed by adding 0.385 ml of 2.5%, 0.77 ml of 5%, 0.77 ml of 7.5%, 0.77 ml of 10%, 0.192 ml of 12.5%, 0.77 ml of 15%, 0.192 ml of 17.5%, 0.192 ml of 20%, and 0.192 ml of 30% iodixanol solutions to the bottom of a 5-ml Beckman centrifuge tube. The 0.7 ml volume gradient was formed by adding 27.5 μl of 30%, 27.5 μl of 20%, 27.5 μl of 17.5%, 110 μl of 15%, 27.5 μl of 12.5%, 110μl of 10%, 110 μl of 7.5%, 110 μl of 5%, and 55 μl of 2.5% iodixanol solutions to the bottom of a 750-μl Beckman centrifuge tube. After centrifugation for 2 h at 150,000g in a SW55i rotor and Beckman Optima centrifuge, 18-25 fractions were collected from the top of the column, depending on the experimental condition. Proteins in the fraction sets were resolved by SDS-PAGE, followed by western blot analyses with antibodies against the proteins of interest, including organelle markers, LARGE, and GluA1. The proteins were quantified by the densitometric analysis of bands in digital images of western blots and subjected to statistical analyses. For each protein, the average density in each fraction was normalized to the peak density of the protein and plotted as a line graph. For organelle markers, each group was initially analyzed separately. However, as no significant changes were observed between wild-type and mutant *Large^myd^* mice, the data were subsequently merged into a single group.

### Immunoprecipitation

Small-scale immunoprecipitation was performed as described previously (Lee et al., 2012), with some modifications.

### Western blot analysis

Western blot analyses were performed as described previously (Kang et al., 2009). The rabbit anti-LARGE antibody (Rb331) (Kanagawa et al., 2004) was characterized previously. The mouse anti-β-actin (Sigma #A5441) and mouse anti-tubulin antibodies (Sigma #T9026) were used at dilutions of 1:10000 and 1:20000, respectively. For organelle markers, antibodies against rabbit anti-GM130 (Abcam #AB52649), mouse anti-TGN38 (Thermo #MA3-063), and mouse anti-P-cadherin (Abcam #AB22744) were used at dilutions of 1:10000. Mouse anti-GluA1 (Millipore #MAB2263), rabbit anti-GluA2/3 (Millipore #07-598), mouse anti-NR1 (Millipore #05-432), and rabbit anti-NMDAR2B antibodies (Abcam #ab65783) were used at dilutions of 1:1000. Quantitative analyses of band intensities were performed using Image Lab (Bio-Rad).

### Protein expression and purification of LARGE and Fc-fused GluA1 Nt, 2Nt, and 4Nt for ELISA

The gene encoding the human LARGE catalytic domain (CD1-CD2) was cloned into a pET21a expression vector (Novagen) using the *NdeI* and *XhoI* restriction sites. The resulting vector was transformed into Origami B (DE3) host cells (Novagen) to facilitate the formation of disulfide bonds. Single colonies were seeded into LB media supplemented with ampicillin (100 μg/ml), kanamycin (50 μg/ml), and tetracycline (15 μg/ml). After an overnight incubation, 10 ml of cultured cells were inoculated into 1,000 ml of fresh LB media. Once the cell density at 600 nm reached approximately 0.5, isopropyl-d-1-thiogalactopyranoside (IPTG) and MnCl2 were added to final concentrations of 0.1 mM and 0.2 mM, respectively, to induce protein expression. The induced cells were further cultured at 18°C for 3 days. Following a cell harvest via centrifugation at 6,000 rpm, the cell pellet was resuspended in lysis buffer (20 mM Tris, 150 mM NaCl, 10 mM imidazole, pH 8.0) and subjected to disruption by sonication. The sonicated lysate was subjected to ultracentrifugation at 13,000 rpm and 4°C for 1 hour, and the supernatant was filtered through a 0.2-μm syringe filter (Millipore) and incubated with His-bind agarose resin (Elpis Biotech, Korea). After washing with a washing buffer (20 mM Tris, 150 mM NaCl, 20 mM imidazole, pH 8.0), LARGE proteins were eluted using an elution buffer containing 200 mM imidazole. Purified LARGE was subjected to a buffer change to a Tris-based buffer (20 mM Tris, 150 mM NaCl, pH 8.0) supplemented with 0.2 mM MnCl_2_. This solution was stored at 4°C for further study.

Fc-GluA1Nt, 2Nt, and 4Nt were purified from HEK293T cells using a transient transfection protocol. Cells were cultured in 10 cm x 15 cm plates to 85-90% confluency and transfected with the target vectors (10 μg/plate). At 72 h post-transfection (Mirus), the cells were harvested via centrifugation at 6,000 rpm, and the cell pellet was resuspended in PBS and lysed by sonication. The lysate was then subjected to ultracentrifugation at 13,000 rpm and 4°C for 1 hour, and the supernatant was filtered through a 0.2-μm syringe filter (Millipore) and incubated with Protein A Sepharose beads (GE Healthcare #17-0780-01). After washing with a washing buffer (0.5% Triton X-100, 0.5 mM EDTA, 0.5 mM in PBS), the target proteins were eluted using an elution buffer containing 0.2 M glycine (pH 2.5). A proteinase inhibitor was added to all purification steps. Purified LARGE was then subjected to a buffer change using a Tris-based buffer (20 mM Tris, 150 mM NaCl, pH 8.0) and stored at 4°C.

### Enzyme-linked immunosorbent assay (ELISA)

A 96-well plate (SPL, Korea) was coated with purified LARGE protein and bovine serum albumin (BSA) at 4°C. The following day, the antigen-coated plate was washed with PBS (pH 7.4) three times, and each well was incubated with a blocking buffer (PBS containing 0.1% Tween-20 and 2% BSA; PBST-BSA) at room temperature for 1 hour. All buffers used in this study were supplemented with 2 mM MnCl_2_. After three washes with PBST, 10 μg/ml of Human-Fc fused GluA1 (GluA1-Fc) was added to the wells and incubated at room temperature for 1 hour. To detect GluA1 binding to the LARGE-coated surface, the plate was washed and incubated for 1 hour with goat anti-Human IgG (Fc specific)-Peroxidase (Sigma #A0170) diluted 1:3,000 in PBST-BSA. Binding signals were developed and stopped by the sequential addition of TMB (3,3',5,5'-tetramethylbenzidine) solution (Sigma #T0440) and 1 N sulfuric acid, and the colorimetric reaction was evaluated by measuring the absorbance at 450 nm. For a competition ELISA, GluA1-Fc was preincubated with 150 μg/ml of soluble LARGE, a competitor, for 30 min before its addition to the LARGE-coated plate. To identify binding preferences of LARGE toward the Fc-GluA1Nt, 2Nt and 4Nt subtypes, different concentrations of GluA1-family proteins were applied to the LARGE-coated plates as described above. The molarity of each subtype was calculated from the band intensity and molecular weight.

### Immunocytochemistry, microscopy, and image analyses

These processes were performed as described previously (Kang et al., 2009), with some modifications.

For most immunocytochemistry experiments, cultured neurons were transfected at day in vitro (DIV) 13 and immunostained after 72 hours. For staining, cells were fixed with 4% paraformaldehyde/4% sucrose in PBS for 15 min and permeabilized in 0.2% Triton X-100 for 10 min at room temperature. After blocking with 10% goat serum, the cells were incubated first with primary antibodies, followed by secondary antibodies. Cell surface proteins were immunostained with antibodies prior to permeabilization.

Immunostained cells were imaged using a confocal microscopy system comprising a Nikon A1 microscope with a 60x oil-immersion objective. The images were analyzed using NIS-Element Software (Nikon). Except for one rabbit anti-GluA1-Ct antibody (this paper: JH4294) and rabbit anti-LARGE antibody (this paper: UT1002), the following primary antibodies were used for staining were obtained commercially: mouse or chicken anti-GFP (Neuromabs #75-131 or Invitrogen #A10262), GluA1-Nt (Millipore), GM130 (BD Transduction Laboratories or Abcam), and chicken anti-MAP2 (Covance #PCK-554P). All secondary antibodies were purchased from Molecular Probes/Invitrogen/Life Technologies.

To quantify the confocal images presented in Figure 3C, the GluA1 intensity in the spine and integrated intensity of individual endogenous GluA1 puncta in the dendritic spine were measured. Images were analyzed using NIS-Elements AR (Nikon). To quantify the confocal images presented in Figure 5C and 5E, fluorescence signals in the soma and dendrites were quantified by measuring the area containing signals above a certain threshold. After normalization, the data were statistically analyzed and plotted as a histogram. To quantify the confocal images presented in Figure 7A-7C, we focused on the number of overlapping Golgi & LARGE, Golgi & GluA1, or LARGE & GluA1 signals. These co-occurrences were independent of signal intensity. Accordingly, we applied the Manders overlap coefficient, which describes the degree of overlap, to the analysis of co-localization. The images were acquired at a thickness of 0.6 μm. The threshold was determined using a global thresholding process to separate pixels from the background, and the coefficient was calculated from the pixels obtained from all slice images. Customized codes written using C++ and MATLAB were used.

### Immunohistochemistry

Following sacrifice, mice were transcardially perfused with solution followed by 4% paraformaldehyde in PBS. The brains were quickly removed and post-fixed in the same solution overnight at 4°C. Coronal sections (50 μm thick) were cut with a vibrotome, washed in PB, permeabilized in PBT. with 0.1% Triton X-100 (Sigma), and blocked with 5% heat-inactivated horse serum (HS) for 1 h. The slices were then incubated with primary mouse anti-GFP (1:500; Neuromab) overnight at 4°C in PB with 5% HS. The slices were subsequently washed in PB and incubated for 2 h at room temperature with an Alexa Fluor 488-conjugated goat anti-mouse antibody (1:1000; Invitrogen). After washing in PB, the slices were mounted using Vectashield with DAPI (Vector #H1200) and stored in the dark at 4°C. High-magnification images were taken using a confocal laser-scanning microscope (Nikon A1 system) equipped with lasers pretuned to 488 nm (i.e., FITC channel) and DAPI. The images were analyzed using NIS-Elements AR (Nikon).

### Quantification and Statistical Analysis

All statistical analyses were performed using SigmaPlot software (Ver 12; SYSTAT Software). The statistical methods used for particular experiments are noted in the figure legends. Each biological experiment was replicated at least three times using different batches of cells or tissues from different animals. The number of required additional experiments was determined using a power analysis, which was based on a statistical analysis of the data from the first three experiments. The final data sets were analyzed using a two-tailed Student’s t-test for experiments with two groups and/or a one (or two)-way ANOVA followed by a *post hoc* Tukey multiple comparison test for experiments with more than two groups. A probability (P) value ≤0.05 was considered significant. All data points were used in plots after confirming a normal distribution (data not shown). Most values are presented as mean values ± standard errors of the means (SEM). Variations were calculated and are presented as SEMs.

## Acknowledgments

This work was supported by the Institute for Basic Science (IBS, R001-D1) (H-S.S.) and the University of Texas Medical Branch (UTMB) (M-G.K.). We thank Drs. Kevin P. Campbell and Richard L. Huganir for the generous gifts of cDNA plasmids, antibodies, and *Large^myd^* mice. We thank all of the Kang Lab members for technical assistance and advice in this study.

## Author Contributions

M-G.K. designed and supervised the research. M-G.K. and B.A.S. wrote the paper. B.A.S., T.C., D.Z.L., H.Y.L., J-J.L. B.L., S-W.K., and M-G.K., performed experiments and analyzed the data. K.A.C., K.T.D., T.A.G., H.M.K., S-Y.C., and H-S.S contributed reagents and analytical tools and provided input and expertise.

---

**Figure 1-figure supplementary 1.**
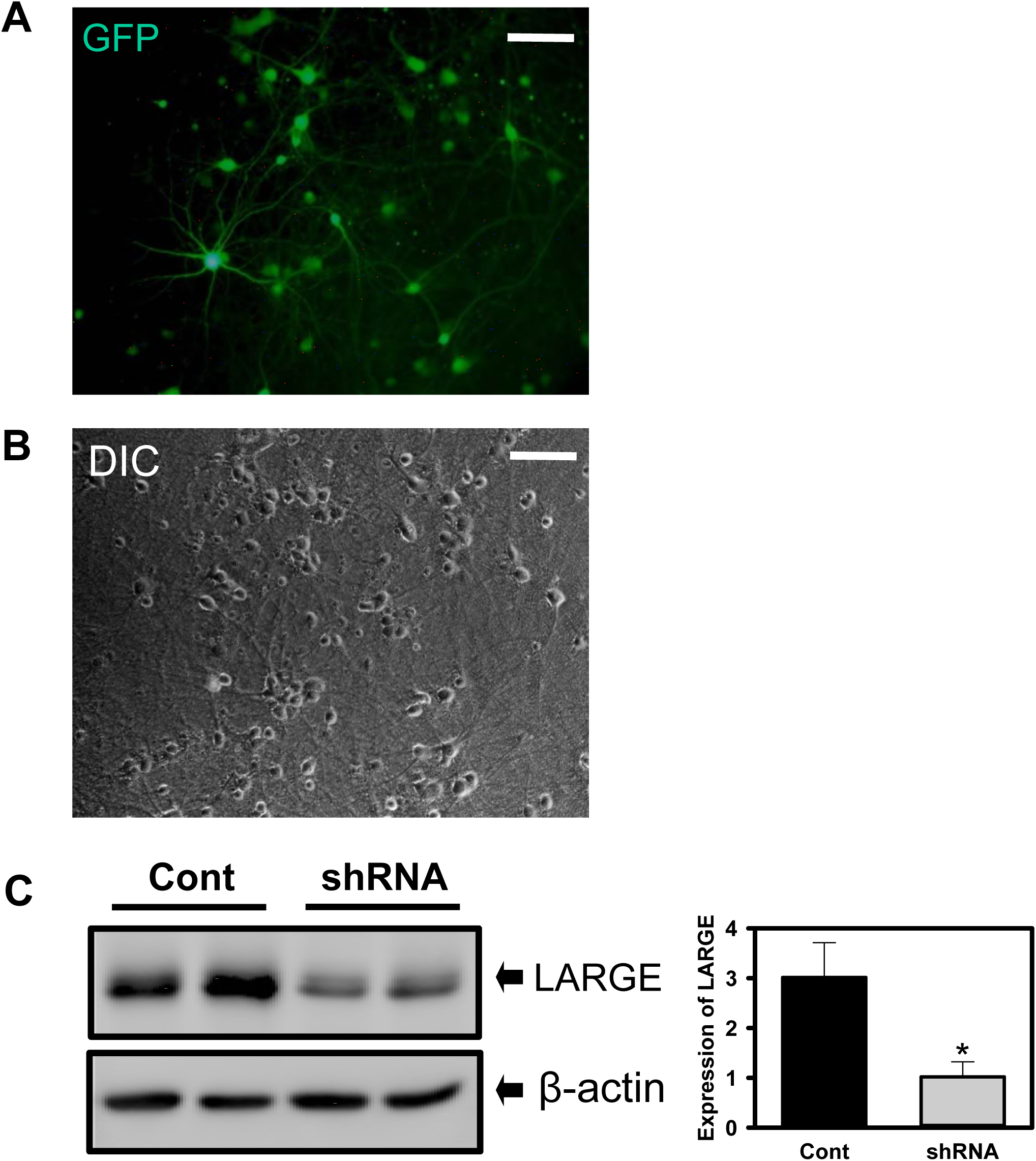
Validation of AAV expressing *LARGE* shRNA with GFP. (**A**) A representative fluorescence image of GFP signals demonstrated that most cultured hippocampal neurons were infected by the AAV. (**B**) Differential interference contrast (DIC) image shows the number and shape of neurons in the culture. These live images were recorded with 20x objective attached to a Nikon TMD inverted microscope system connected to a Nikon digital camera system (Digital-Sight DS-2Mv). (**C**) Western blot analysis confirmed knockdown of *LARGE.* Scrambled shRNA was used as control (Cont) (n=3, two tailed t-test, *\*P* < 0.005). Scale bar = 100 μm.

**Figure 1-figure supplementary 2.**
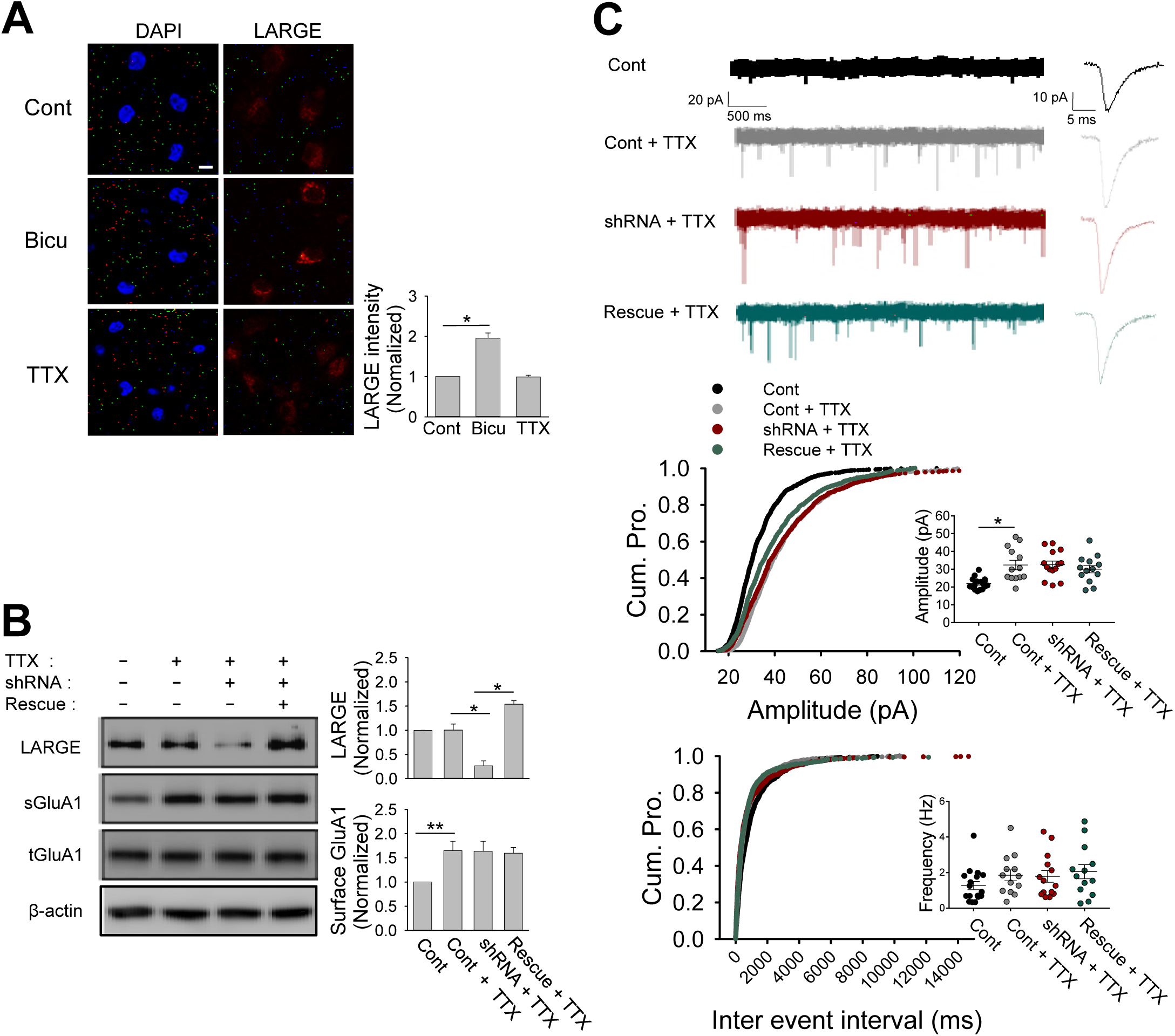
LARGE is necessary for neuronal homeostatic scaling-down but not for scaling-up. (**A**) Confocal images of cultured hippocampal neurons confirmed the increase in LARGE expression in response to Bicu but not TTX (n = 13, 13, 13 neurons; Mann-Whitney rank sum test, T = 91, U = 0, \**P* <0.001; Blue: DAPI, red: endogenous LARGE). Scale bar = 10 μm. (**B**) The TTX-induced increase in surface GluA1 was not affected by LARGE shRNA or LARGE rescue. (n = 3; one-way ANOVA, \**P* <0.001, \*\**P* = 0.027). (**C**) The TTX-induced increase in mEPSC amplitude was not affected by LARGE KD or rescue (1300, 1270, 1300, and 1279 events from n = 17, 13, 14, and 13 neurons, respectively; one-way ANOVA; amplitude, F(3,53) =7.783, \**P* <0.001; frequency; F(3,53) =1.211, P = 0.315). The mEPSC traces and cumulative and scattered plots are shown. Nine different groups were recorded using 60 coverslips from three batches of neuronal culture.

**Figure 1-figure supplementary 3.**
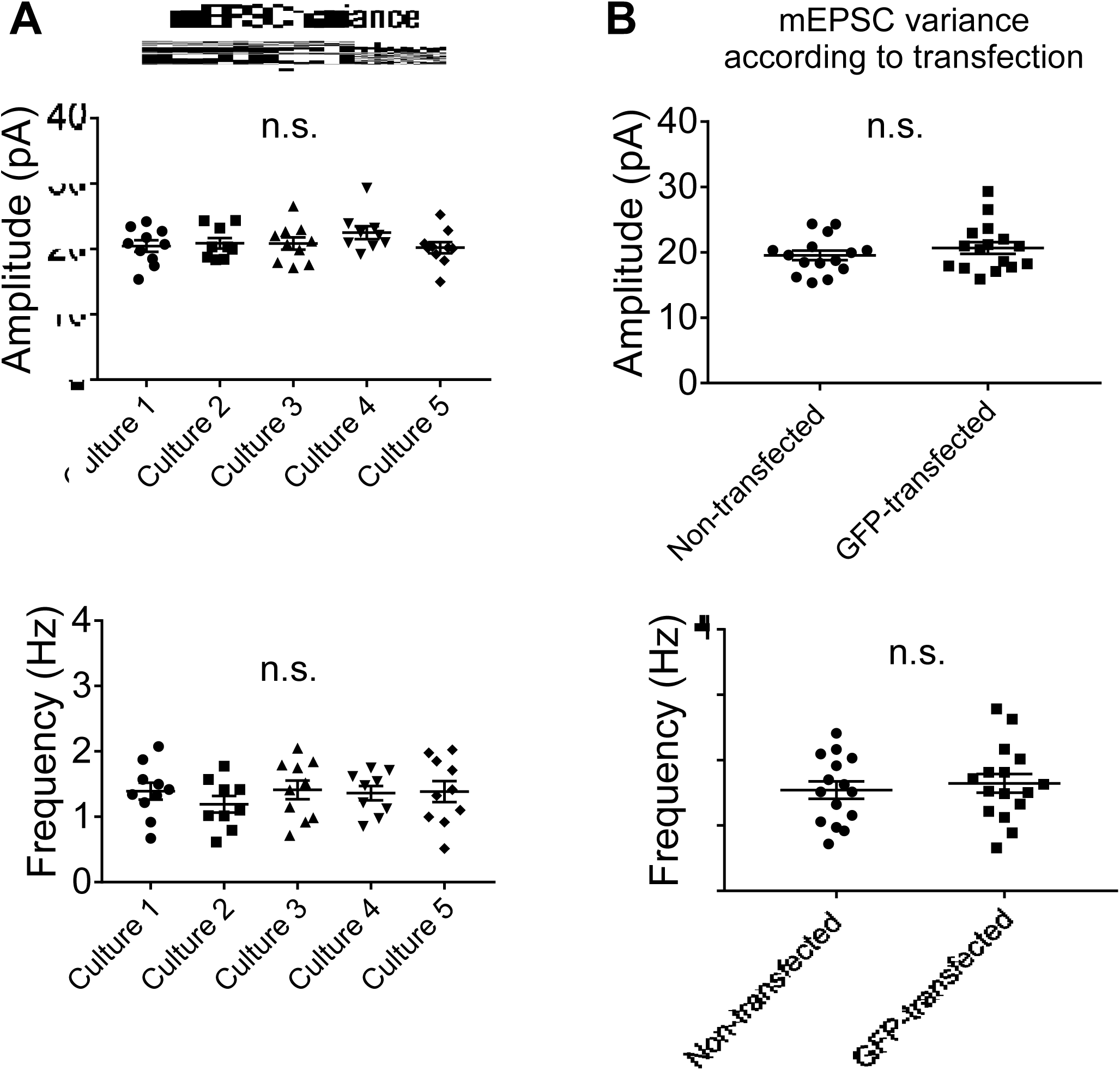
Verification of low-level batch-to-batch variation in our mEPSC experiments. (**A**) The amplitude and frequency did not vary among the batches used in experiments (**B**) The amplitude and frequency did not vary between non-transfected and GFP-transfected neurons

**Figure 3-figure supplementary 1.**
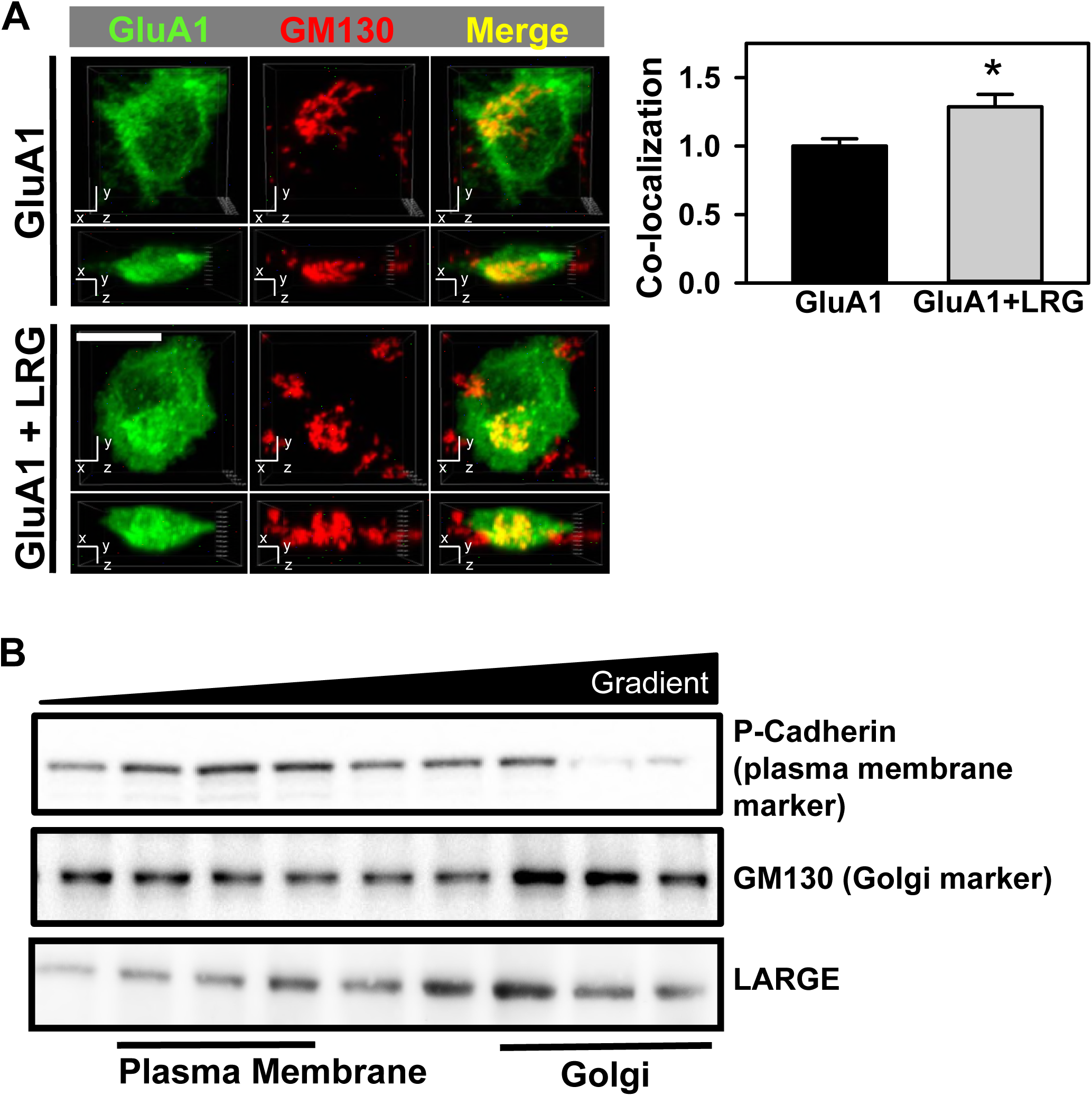
LARGE increased AMPA-R pool in the Golgi. (**A**) GFP-GluA1 in Golgi was increased by LARGE overexpression (GluA1 + LRG), compared with a control (GluA1) (n = 30, 30 cells; Two tailed t-test, \**P* < 0.001). Colocalization of GluA1 with GM130 in HEK293T cells analyzed by 3D reconstruction of a series of z-stack confocal images. Complete (yellow) and partial co-localization (orange). Co-localization of GluA1 and LARGE was quantified using the colocalization analysis tool in the NIS-Elements software (Nikon). In the analysis, Manders overlap coefficients were given and used to obtain the relative colocalization values between those two proteins. Scale bar = 10 μm. (**B**) Fractionation of subcellular organelles from the hippocampal CA1 of mice. Representative data showing the fractionation of organelle markers and LARGE. P-cadherin, plasma membrane marker; GM130, Golgi markers.

**Figure 3-figure supplementary 2.**
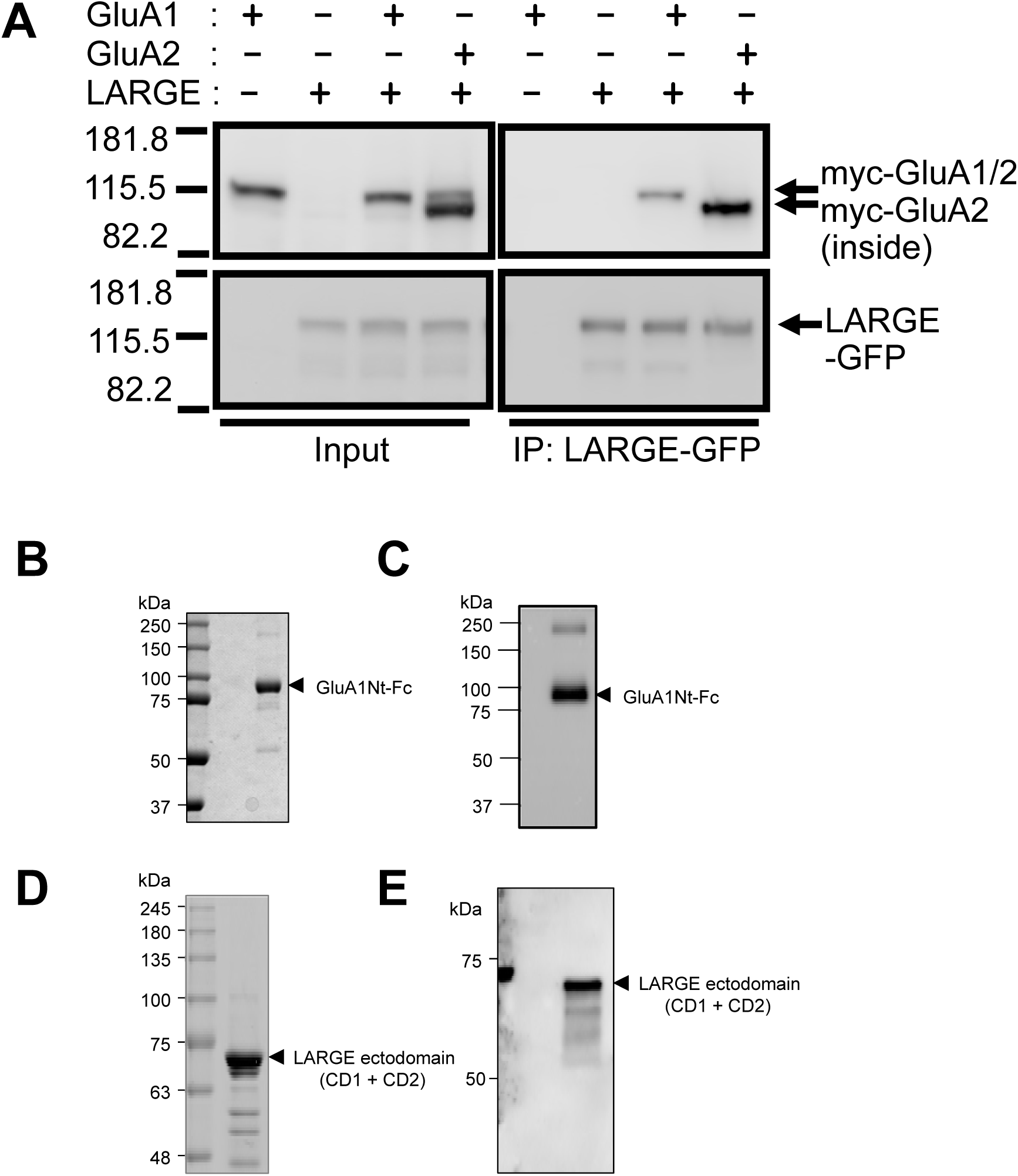
LARGE directly interacts with AMPA-R subunits. (**A**) LARGE interacted with GluA2 inside of cell (lower band) but not with GluA2 at the plasma membrane (upper band). Analysis of LARGE association with AMPA-R determined by immunoprecipitation (IP). HEK293T cells were transfected with myc-GluA1, myc-GluA2, and/or LARGE-GFP. IP with anti-GFP antibody. Both GluA1 and GluA2 were co-immunoprecipitated with LARGE-GFP. As see in Input, GluA2 yield two bands (upper and lower bands). Most GluA2 co-immunoprecipitated with LARGE was GluA2 correspond to intracellular GluA2, judging from its molecular weight. (**B-E**) Purification of GluA1 and LARGE (catalytic domain 1 [CD1] + CD2) proteins. SDS-PAGE (**B**) and Western-blot (**C**) analysis of purified GluA1Nt-Fc fusion protein. SDS-PAGE (**D**) and Western-blot (**E**) analysis of purified LARGE.

**Figure 3-figure supplementary 3.**
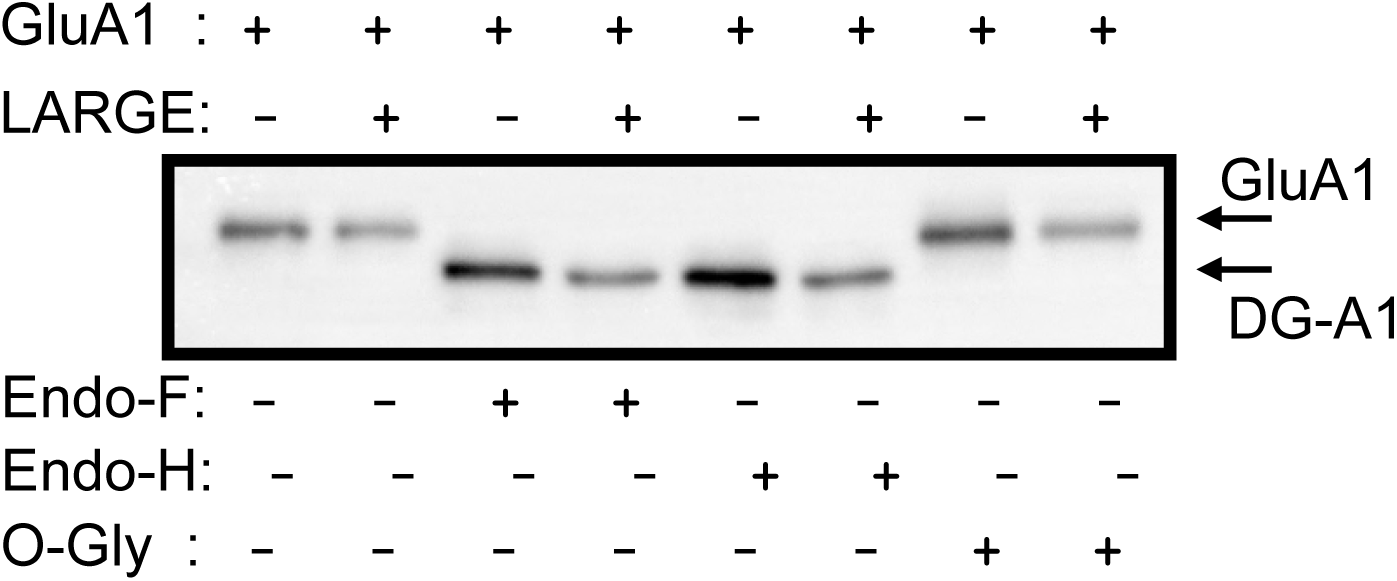
LARGE interaction with AMPA-R did not change glycosylation of AMPA-R. No change in N- and O-glycosylation of GluA1 with coexpression of LARGE in HEK293T cells. N-linked glycosylation of GluA1 was analyzed using two de-glycosylation enzymes, endoglycosidase F (Endo-F) and endoglycosidase H (Endo-H). Endo-F completely de-glycosylated all GluA1 regardless of LARGE co-expression, and there was no difference in the amount of Endo-H-sensitive and -insensitive forms of GluA1 with co-expression of LARGE. In addition to N-linked glycosylation of GluA1, O-linked glycosylation of GluA1 was analyzed using O-glycosidase (O-Gly). The size of a GluA1 band was not changed by O-Gly treatment, suggesting that GluA1 does not have O-glycosylation. Coexpression of LARGE did not changed the O-glycosylation status of GluA1. Method: In HEK293T cells, myc-GluA1 and LARGE were expressed by transfection of their plasmids. From the lysate of the cells, myc-GluA1 was immunoprecipitated and treated with Endo-F (500 unit) and Endo H (500 unit) at 37 °C for 2 hours, or with O-Gly (50000 unit) followed with SDS-PAGE and Western blot analyses.

**Figure 5-figure Supplementary 1.**
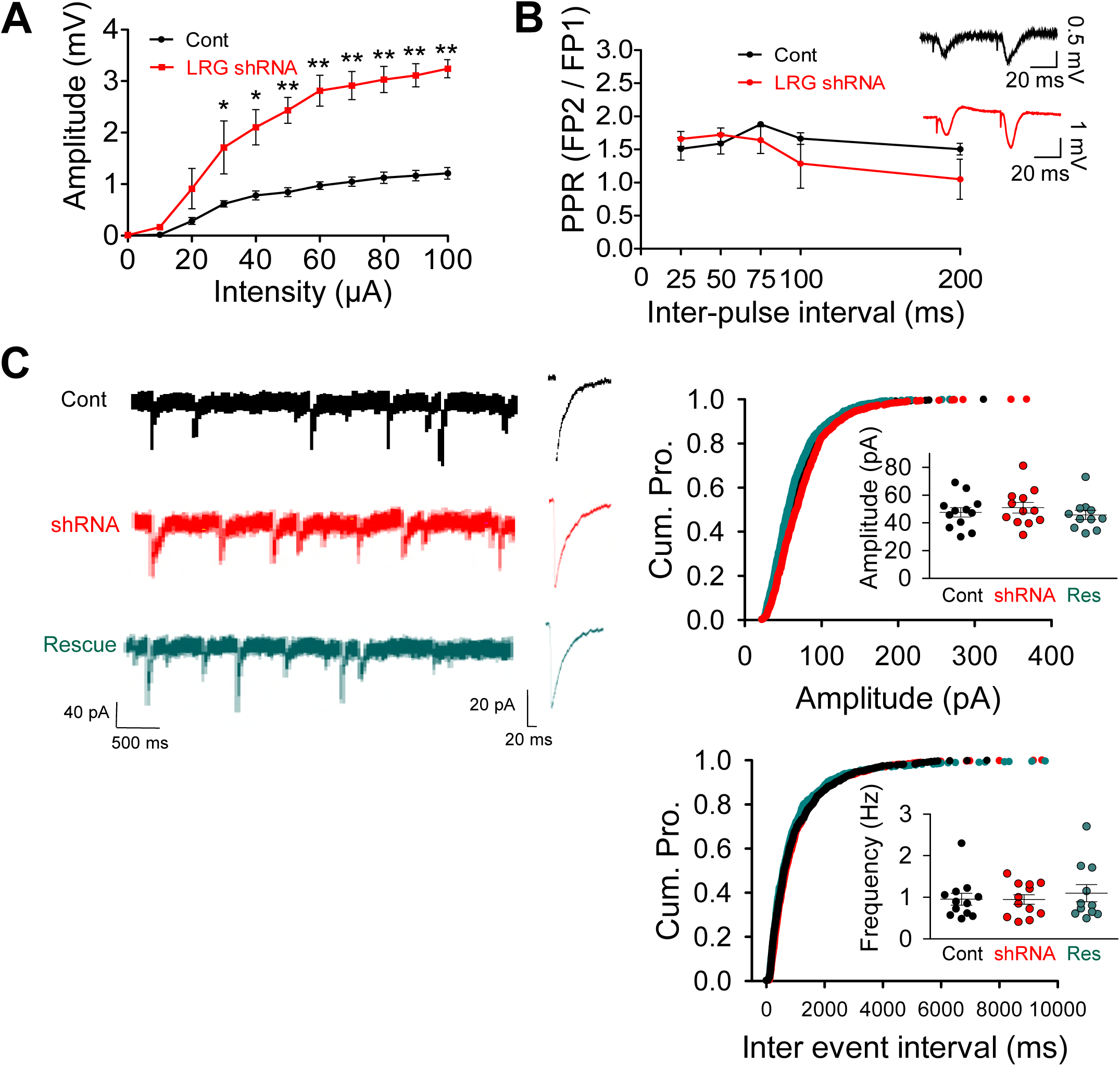
Electrophysiological analyses after knockdown of *LARGE* at CA1 of hippocampus. (**A**) AMPA-R fEPSP amplitudes were significantly increased by LARGE KD. Input-output curves shown relating stimulus strength to fEPSPs (output amplitude), which is greater in a LRG shRNA group *(n =* 5, 5 mice; Two-way repeated-measures ANOVA with post hoc Bonferroni t-test, *\*P* < 0.05, \*\**P* < 0.001). (**B**) Intact short-term plasticity. Mean PPR (FP2/FP1) of fEPSPs plotted as a function of inter-pulse. Field potential (FP) data are mean ± s.e.m. (group: *P* = 0.565, group x interval: *P* = 0.170, n=5, 5). (**C**) LARGE did not affect inhibitory synaptic strength. mIPSC recordings from Cont, shRNA, and Rescue. No significant differences in mIPSC amplitude and frequency were observed among the groups. Three different groups were recorded using 27 coverslips in 3 batches of neuronal culture (1098, 1080, 1055 events from n = 12, n=12, n=11 neurons, respectively; Amplitude, F(2,32)=0.553, *P* =0.58; Frequency, F(2,32)=0.292, P=0.749). Scale bar = 30 μm.

**Figure 5-figure supplementary 2.**
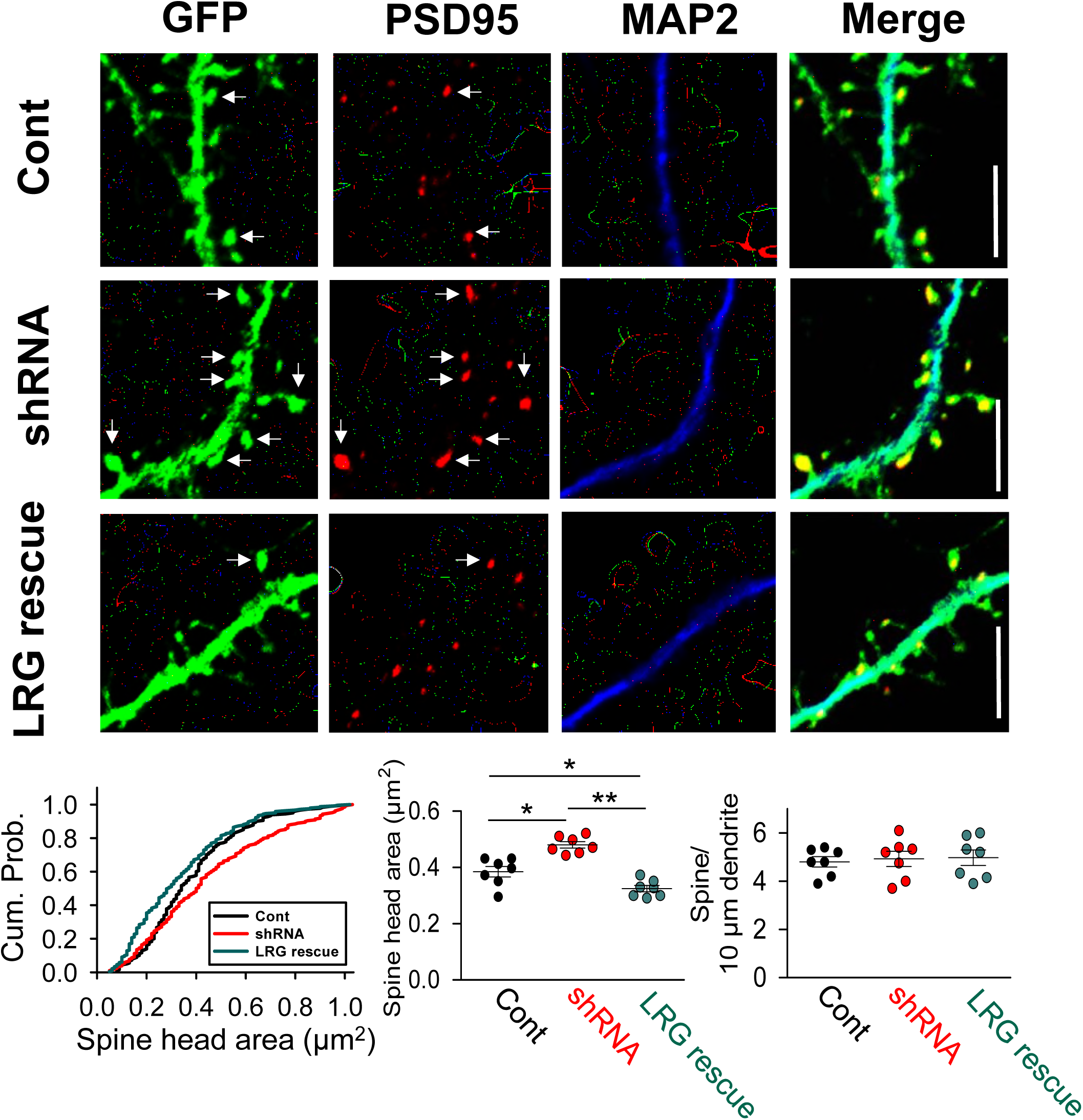
Confocal image analyses demonstrated the effect of *LARGE* KD on structural synaptic plasticity. Confocal images of cultured hippocampal neurons showed that *LARGE* KD (shRNA) significantly increase not the number of spines but the size of spines compared to that of control neurons, which was reversed by LARGE rescue *(n =* 7, 7, 7 neurons; One-way ANOVA, \**P* < 0.01, \*\**P* < 0.005). In the cultured hippocampal neurons, neuronal dendrites and spines was visualized by GFP. PSD95 is synaptic marker. MAP2 is dendrite marker. Scale bar = 10μm.

**Figure 6-figure supplementary 1.**
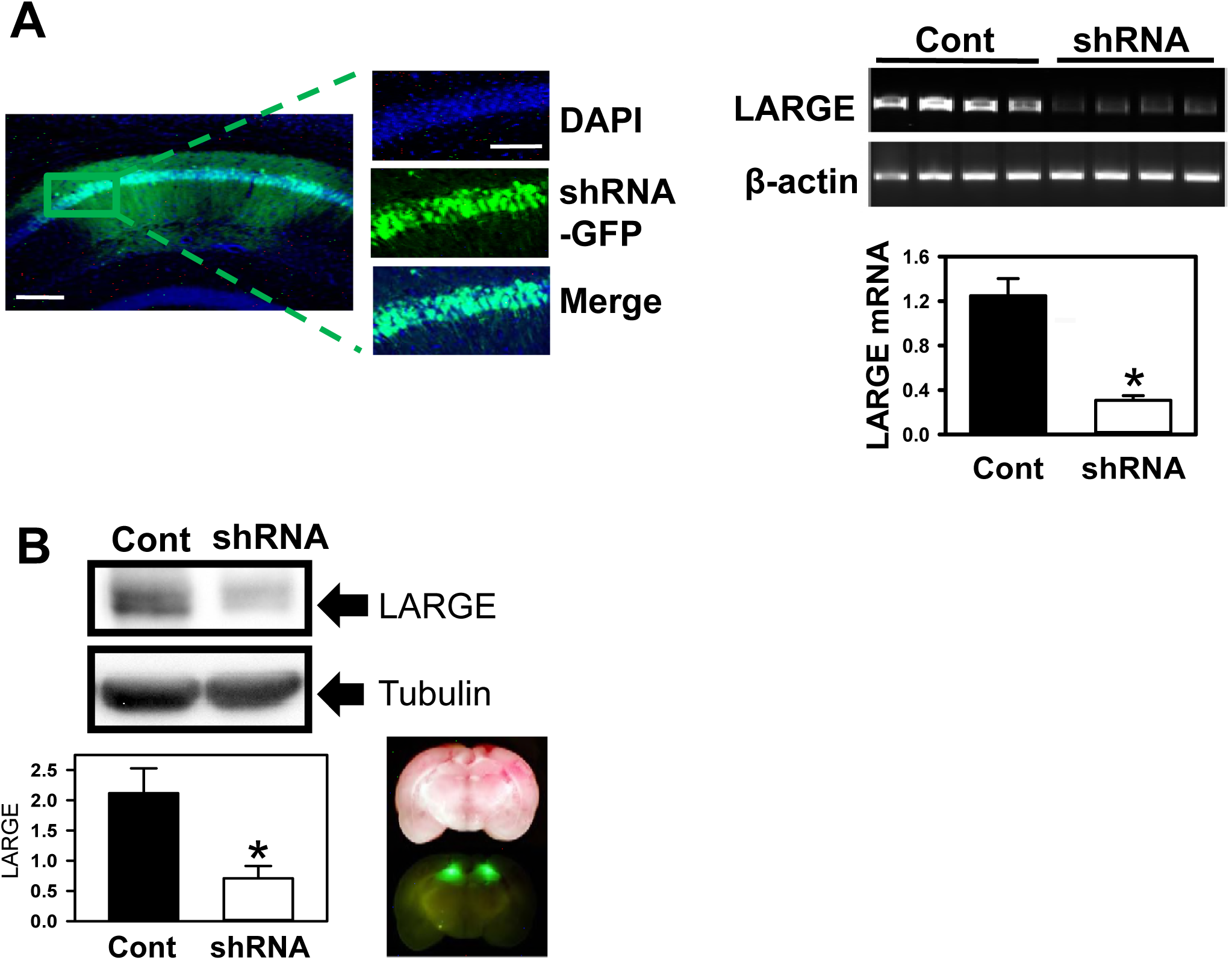
Confirmation of knockdown efficiency of LARGE shRNA after *in vivo* electrophysiology analyses and behavior tests. (**A**) Immunohistochemistry and RT-PCR analysis of mouse hippocampal CA1 region infected with AAV. Confocal images showed the location and diffusion range of AAV microinjected into CA1. Endogenous *LARGE* mRNA expression in CA1 was significantly knocked down by infection of AAV expressing *LARGE* shRNA with GFP (shRNA) (n = 3, 3 mice; Two tailed t-test, \**P* < 0.001). Scale bar = 100 μm (left), 50 μm (right). (**B**) Digital image and Western blot analysis of hippocampal CA1 region of rat brain infected with AAV. Brain slices were imaged by digital imaging under a blue LED light. Strong and specific expression of GFP in hippocampi indicated specific delivery and expression of *LARGE* shRNA with GFP by AAV injection. Western blot analyses confirmed knockdown of *LARGE* in GFP-expressing hippocampi *(n =* 3, 3 rats; Two tailed t-test, \**P* < 0.005)

**Figure 6-figure supplementary 2.**
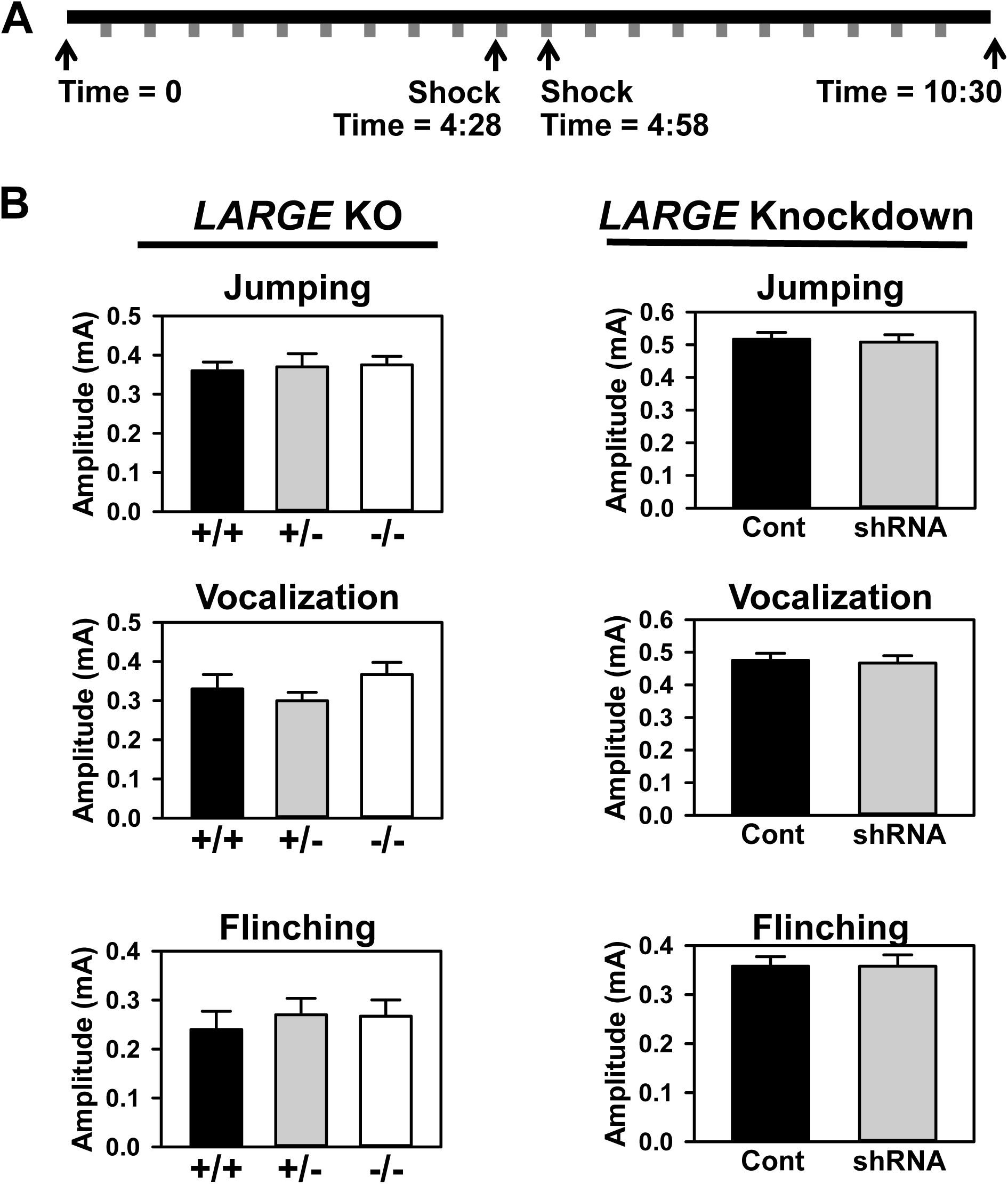
The absence of a significant difference in shock threshold among groups demonstrated that LARGE expression status did not affect fear conditioning. (**A**) Schema of shock threshold tests of animals. Shocks (gray blocks) were delivered every 30 s, with intensity increasing from 0.1 mA to 1.0 mA (4:28, 4:58) and decreasing back to 0.1 mA (10:30). (**B**) Regardless of genotype or injected AAV, all animals responded to shocks in a similar way. The shock intensity thresholds for jumping, vocalization, and flinching were measured for wild type (+/+), heterozygous (+/-), and knockout (-/-) animals and animals injected with AAV expressing scrambled shRNA with GFP (Cont) or *LARGE* shRNA with GFP (shRNA).

